# Spatio-temporal “global” neurodynamics of the human brain in continuous and discrete picture: Simple statistics meet on-manifold microstates as multi-level cortical attractors

**DOI:** 10.1101/2023.07.13.548951

**Authors:** Tomohisa Asai, Shiho Kashihara, Shinya Chiyohara, Kentaro Hiromitsu, Hiroshi Imamizu

**Affiliations:** Advanced Telecomunications Research Institute International (ATR); The University of Tokyo

**Keywords:** Microstate, Neural manifold, state space, EEG, fMRI, MEG, mass global dynamics

## Abstract

The neural manifold in state space represents the mass neural dynamics of a biological system. A challenging modern approach treats the brain as a whole in terms of the interaction between the agent and the world. Therefore, we need to develop a method for this global neural workspace. The current study aimed to visualize spontaneous neural trajectories regardless of their measuring modalities (electroencephalography [EEG], functional magnetic resonance imaging [fMRI], and magnetoencephalography [MEG]). First, we examined the possible visualization of EEG manifolds. These results suggest that a spherical surface can be clearly observed within the spatial similarity space where canonical microstates are on-manifold. Once valid (e.g., differentiable) and useful (e.g., low-dimensional) manifolds are obtained, the nature of the sphere, such as shape and size, becomes a possible target of interest. Because these should be practically useful, we suggest advantages of the EEG manifold (essentially continuous) or the state transition matrix (coarse-grained discrete). Finally, because our basic procedure is modality-independent, MEG and fMRI manifolds were also compared. These results strongly suggest the need to update our understanding of neural mass representations to include robust “global” dynamics.

## Introduction

Generally, the neural manifold represents the state space for mass neural activity as a system in general (Iyer et al., 2022; Jazayeri & Afraz, 2017). A traditional approach spatiotemporally divides the brain into parcels or modules in relation to a specific perceptual or cognitive function. In addition, the integration of segregated modules into a global neural workspace is important for understanding brain function. A challenging modern approach would be to treat the brain as a whole in terms of the interaction between the agent and the world (Friston et al., 2021; Mathys et al., 2011) in which the essential function of the brain is simply the interpretation of sensory inflow and motor outflow over multiple (hierarchical) representational spaces (Barack & Krakauer, 2021; Kriegeskorte & Wei, 2021). In this view, whole-brain dynamics as an agent of the entity would need to be visualized for both its common and unique features among participants. While previous studies have often examined neural manifolds for a relatively local population of neurons (Nieh et al., 2021; Sadtler et al., 2014), others have indicated more global neural dynamics as a whole-brain state transition both in functional magnetic resonance imaging (fMRI) (state-space landscape) (Ezaki et al., 2017; Watanabe et al., 2014) and electroencephalography (EEG microstate) (Lehmann, 1971; Pascual-Marqui et al., 1995). This strategy produces a state vector at each moment within the phase space (Jazayeri & Afraz, 2017; Ros et al., 2014; Shaw et al., 2019). Tracking the trajectory of the state vector (i.e., a point) further embodies the neural manifolds as an assembly. The current study aimed to visualize spontaneous neural trajectories regardless of the measuring modality (EEG, fMRI, or magnetoencephalography [MEG]) to provide further insight into neural mass representation, given its robust global dynamics.

### EEG canonical microstates as attractors

The current approach is motivated by EEG microstate (EEGms) analysis. EEGms have been robustly reported as quasi-stable potentials lasting 60–120 milliseconds that represent whole-brain state dynamics (i.e., network activity) (Michel & Koenig, 2018). The common procedure for EEGms analysis comprises two stages. At the *clustering* stage, the fluctuation in the standard deviation among all electrodes represents the global field power (GFP) for each recording. The local maxima of the GFP (i.e., peaks) are assumed to best represent periods of momentary stability as the state in the voltage topography (Zanesco, 2020), while local minima are assumed to be transient moments between the states. For example, imagine a chain of mountains as a landscape in which each mountain represents a unique phase. The peak is the “limit” representation of the mountain, while the bottom is between the mountains. Therefore, every spatial pattern at the GFP peaks was accumulated for the participant group (the GFP peak dataset). An unsupervised learning algorithm, such as K-means, clusters the GFP peak dataset into optimal classes. Finally, group-wise common topographies are obtained as templates for each class (typically msA, B, C, and D). This clustering procedure aims to extract a few “attractors” in the state dynamics (Férat et al., 2021; Milz et al., 2017) that are often expressed as the cluster centroids on an abstract space (possibly high dimensional) (Koenig & Brandeis, 2016; Zanesco, 2020). Therefore, the continuous spatiotemporal dynamics of the EEG state can be depicted as a transition among these attractors (four canonical EEGms in the current case)(Mishra et al., 2020). This coarse-graining discretization produces a state-transition matrix or directed graph (consisting of nodes and edges) through the *labeling* stage.

The labeling procedure measures the similarity between the templates obtained and every time point (including the GFP peak points) of the participants and labels all data points as one of them (e.g., msA) in a “winner-take-all” manner (Mishra et al., 2020; Pascual-Marqui et al., 1995). Because each topography should last for a while on the order of milliseconds, the original EEG data with multiple channels are simply visualized as a state transition pattern among four states, such as C, D, B, A, and D. As a result, the EEGms analysis typically reveals the frequency of the state (“occurrence”), the lasting of the state (“duration”), and transition patterns among the states (“transition probability”) for each recording session or participant.

### Continuous or discrete

These depicted features of EEGms can be compared among participant groups (e.g., patients with a disorder and controls) as potential biomarkers (de Bock et al., 2020; Perrottelli et al., 2021) because this discretization of dynamics is relatively stable, regardless of some variations in the analysis algorithms (Wegner et al., 2018). However, these features can sometimes go alone and overemphasize the static traits, that is, the state transition might be assumed as the actual depiction of discrete EEG state dynamics, possibly with “jumping” (Mishra et al., 2020; Shaw et al., 2019). Although EEGms analysis and calculated features are useful tools, especially for comparing people in a simpler manner, we need to consider the possible continuous representation of EEG state dynamics as a whole, that is, the neural and statistical manifolds within the phase (state) space. This approach would ensure detailed characterization of the atypical EEG dynamics for a specific population. In this sense, continuous-or-discrete is not the opposite but the tandem approach.

### The current study

First, we examined the possible visualization of EEG manifolds from a simple thought experiment to actual manifold learning. Once valid (e.g., differentiable) and useful (e.g., low-dimensional) manifolds are obtained, the nature of the manifold, including shape and size, becomes a possible target of interest. Furthermore, if we are interested in group-wise dynamics in the EEG state, hyper alignment across participants is required to standardize the individual manifold. Because these should be practically useful, we suggest advantages of the EEG manifold (essentially continuous) or the state transition matrix (coarse-grained discrete). Finally, because our basic procedure is modality-independent, MEG and fMRI manifolds were also compared. Based on these results, we seek to find the neural trajectory as analogous to the trajectory of body state (e.g. hand positions and joint angles) where both are moving toward an agent-dependent “meaningful position” within a certain space: the former represents phase-shifting to a task-specific neural state, while the latter practically means to reach to a cup.

## Materials and Methods

The following results were obtained from several datasets, including our own multiple data (different EEG caps, MRI scanners, subjects, and experiments) and some other external open datasets (e.g., Babayan et al., 2019; Niso et al., 2016). Although we do not specifically mention the source for each result, we have confirmed that the depicted results are not dependent on the data, regardless of the measuring procedure, devices, subjects, and preprocessing policy.

## Results and Discussion

### Characteristics of the individual manifolds

The traditional understanding of EEGms is the momentary state transition among the four typical topologies (spatial representations as canonical msA, B, C, and D). These simplified features for neurophysiological dynamics often help to predict other cognitive or mental variables (da Cruz et al., 2020; Michel & Koenig, 2018)). In contrast, the EEGms transition could simply be a discretization over spatiotemporally continuous dynamics (Mishra et al., 2020). If a “smooth trajectory” (i.e., no jumping) within a certain state space is observed (Shaw et al., 2019), a “smooth surface” as a neural manifold might also emerge where canonical templates serve as a basis as Figure 1A suggests (Fig. 1B for polarized assumption). The term “manifold” has been used to imply relatively low-dimensional and smooth dynamics (Iyer et al., 2022). For that purpose, we first examined the intrinsic dimensionality of the EEG channel space by applying principal component analysis. Figure 1C suggests that the three dimensions are intuitively sufficient to represent global state dynamics because the cumulative proportion of variance almost equals or exceeds 80% at the third PC. To visualize a latent manifold, possible dimensional reductions (e.g., Moon et al., 2019 for more general biological data) from the original channel space (in this case, 32D from 32 electrodes) to a sufficiently low-dimensional space (e.g., 3D in contrast to 2D) were examined (Fig. 1D). Although we may see method-dependent state spaces respectively, a “manifold” in a more strict sense (i.e., smooth or “geometrically differentiable” surface) was only observed within the similarity space by a multi-dimensional scaling (MDS)(e.g., Kriegeskorte & Wei, 2021). Interestingly, this simple spherical surface represented all possible EEG states using a standard MDS procedure (Fig. 2) (see also Supplementary Animation S1).

**Figure 1.**
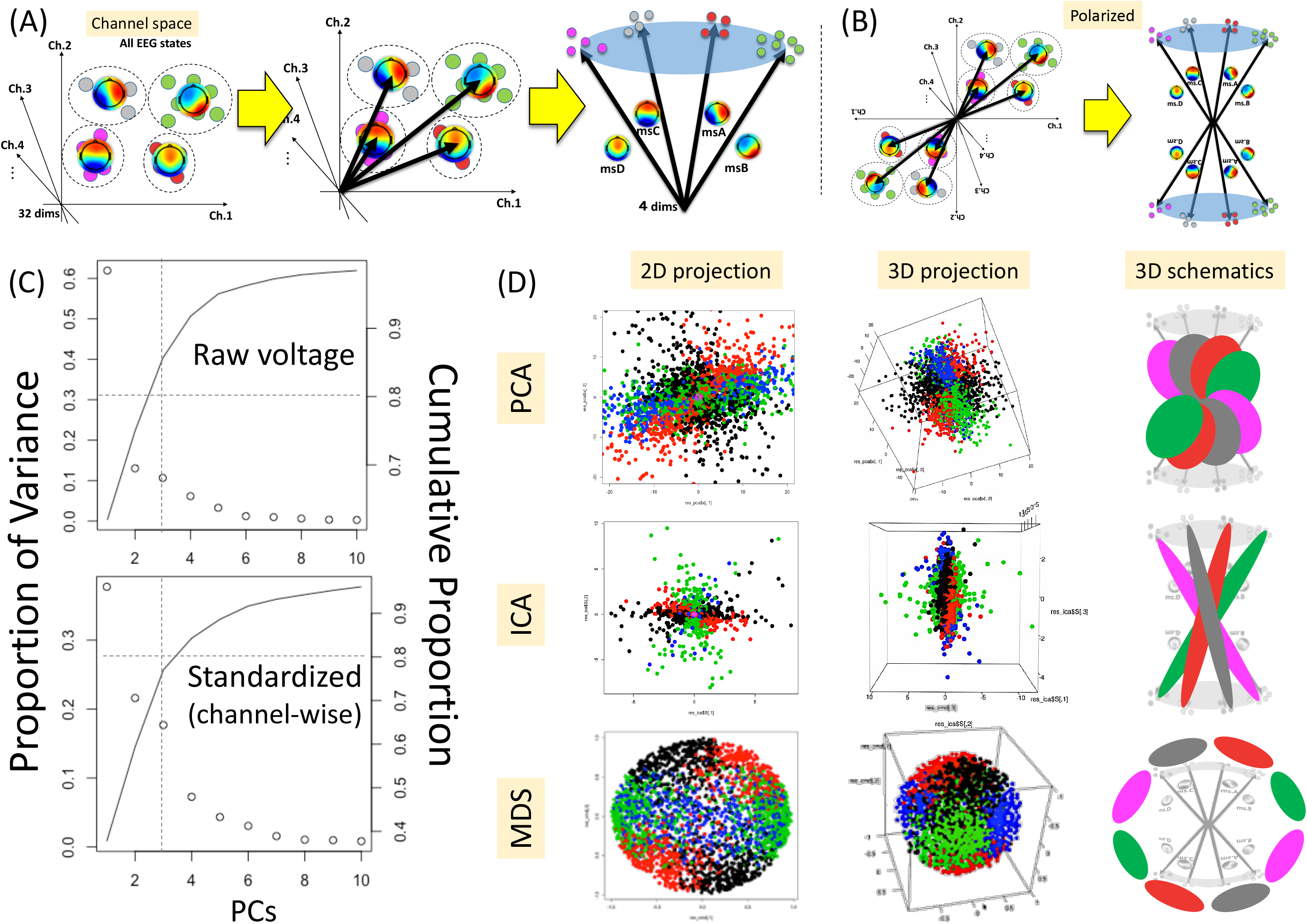
Visualizing manifolds

**Figure 2.**
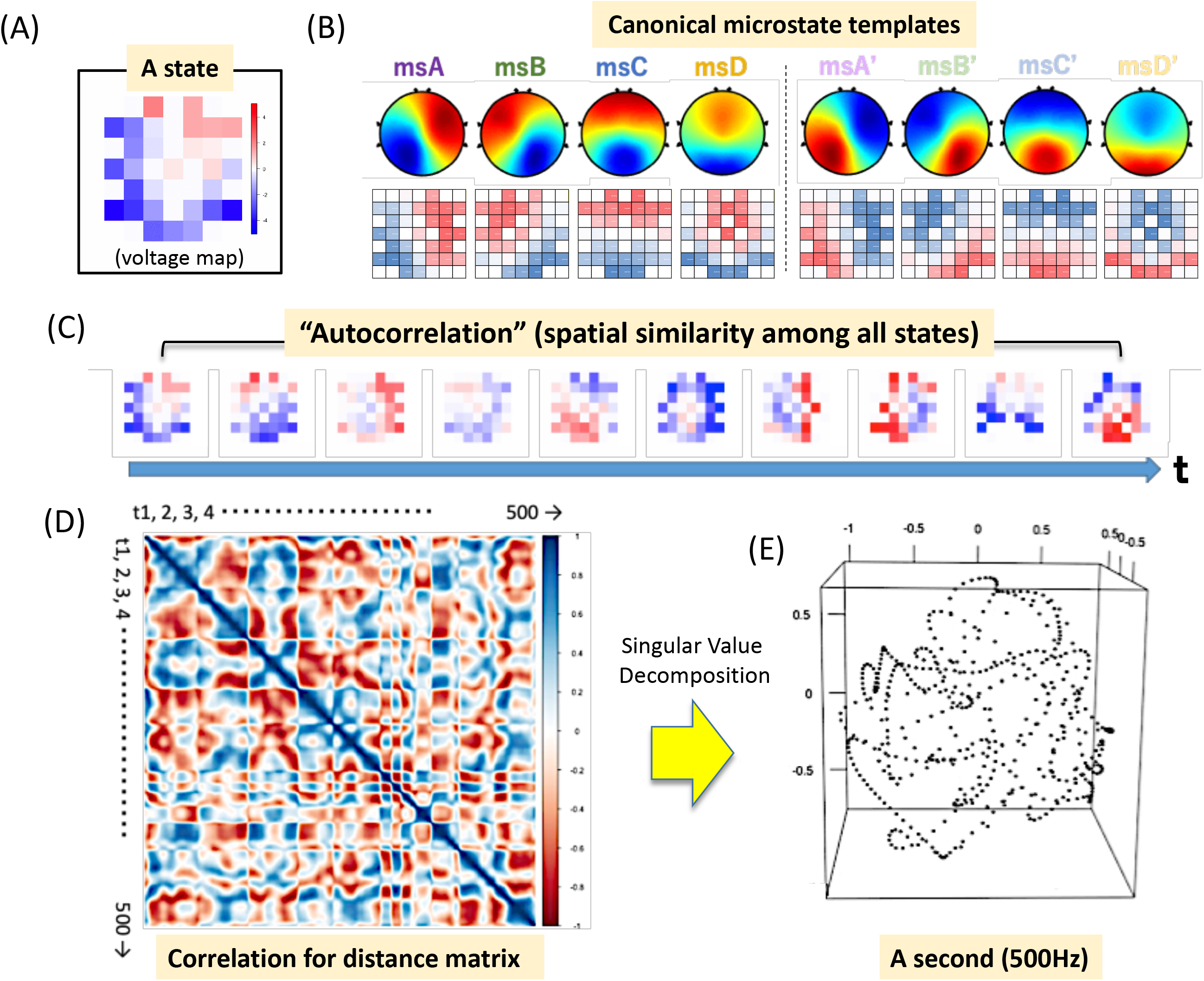
Standard MDS procedure

The state comprises elements at a given moment in the definition (Fig. 2A). Because the elements (32 channel voltage values in the current case) are vectorized, the assembly of states can be seen as a matrix (32 × number of states). As previously mentioned, the conventional EEGms analysis pipeline extracts templates from this matrix (specifically, the GFP peak dataset). The resultant cluster centroids were restored to their original channel configuration (Fig. 2B, upper panel with spatial interpolation from the lower panel). Since the modified k-means was introduced to produce polarity-ignored EEGms templates (Pascual-Marqui et al., 1995), the produced maps can be reversed for polarity-assumed templates, where the frontal is red for positive templates and blue for negative templates (Asai et al., 2022). In addition to EEGms, the assembly of states as a matrix can be time-series EEG data (Fig. 2C; for example, 500 time points). We can calculate the spatial correlation among all states for a similarity score as correlation matrix ranging from -1 to 1 (Fig. 2D, 500 × 500, in this case). If we see something in its patterns (e.g., nonlinear waves or “ripples”), the time-series dynamics as a trajectory might create a closed shape as a manifold. The MDS algorithm dissolves the distance matrix (converted from the correlation matrix) into xyz positional configurations for all states using singular value decomposition. As a result, the exemplified 500 points (for 1 s at 500 Hz sampling) in this state space are *temporally* continuous (Fig. 2E). Furthermore, since this state space is defined as a spatially similarity space, a temporally continuous trajectory within this also means *spatio-temporally* continuity (Fig. 3C, Supplementary Animation S1).

**Figure 3.**
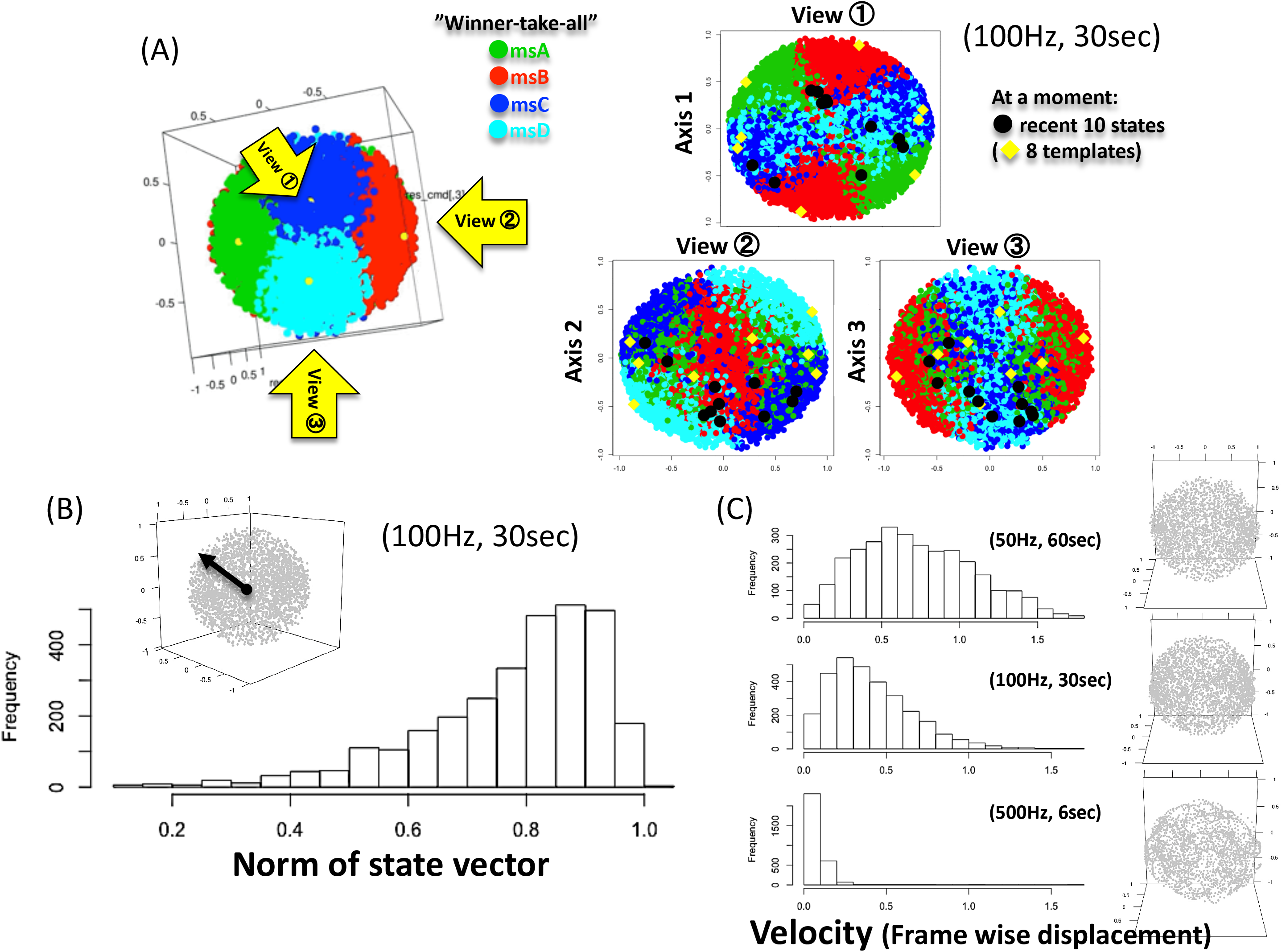
Spherical EEG manifolds

Finally, when we gathered as many states as possible (e.g., for 30 s), the neural manifold emerged as a sphere. The canonical EEGms templates with polarity (Fig. 2B) are located on the sphere surface (Fig. 3A), where each state is colored in a winner-takes-all manner (left, Fig. 5) or where the current state moves on this manifold (right). The norms of the state vectors within this space are distributed toward almost 1 (Fig. 3B), such that the spherical surface, whose radius is almost 1, could be a definition of a whole-brain EEG manifold. When we visually confirmed the actual movement of the state on this manifold, it was clearly temporally continuous (i.e., no jumps) (c.f., Shaw et al., 2019), especially when the sampling rate was high (Fig. 3C). If the sampling rate is sufficiently high for EEG dynamics, there should be no large displacements from the previous time point if we assume spatiotemporally continuous dynamics. The comparison in the framewise displacement among the distributions for different sampling rates indicates that is the case (Fig. 3C upper) where the displacement “1.5” or “2,” for example, means the jump to antipodal point in the current case (e.g., from Japan to Brazil).

Because the canonical templates indicate attractor states (Milz et al., 2017) or attractor position in our case, given a typical EEGms analysis for the GFP peak dataset, we expected a relationship among GFPs, distances to templates, and norms of state vectors (Koenig & Brandeis, 2016; Zanesco, 2020). Although GFP (or log GFP) was non-linearly correlated with the norms of state vectors (Fig. 4A), this positive relationship indicates that when the GFP of the state is higher, the state is located nearer to the spherical surface within the MDS space (and is located nearer to the templates in the definition because templates are located on the surface; Fig. 3A). When each state was colored in a winner-take-all manner, msC was the strongest attractor among the four with the highest GFPs (rightward in Fig. 4BC) (Bagdasarov et al., 2022; Zanesco et al., 2020) whereas msD was concentrated on the sphere surface with the largest norms (upward in Fig. 4B).

**Figure 4.**
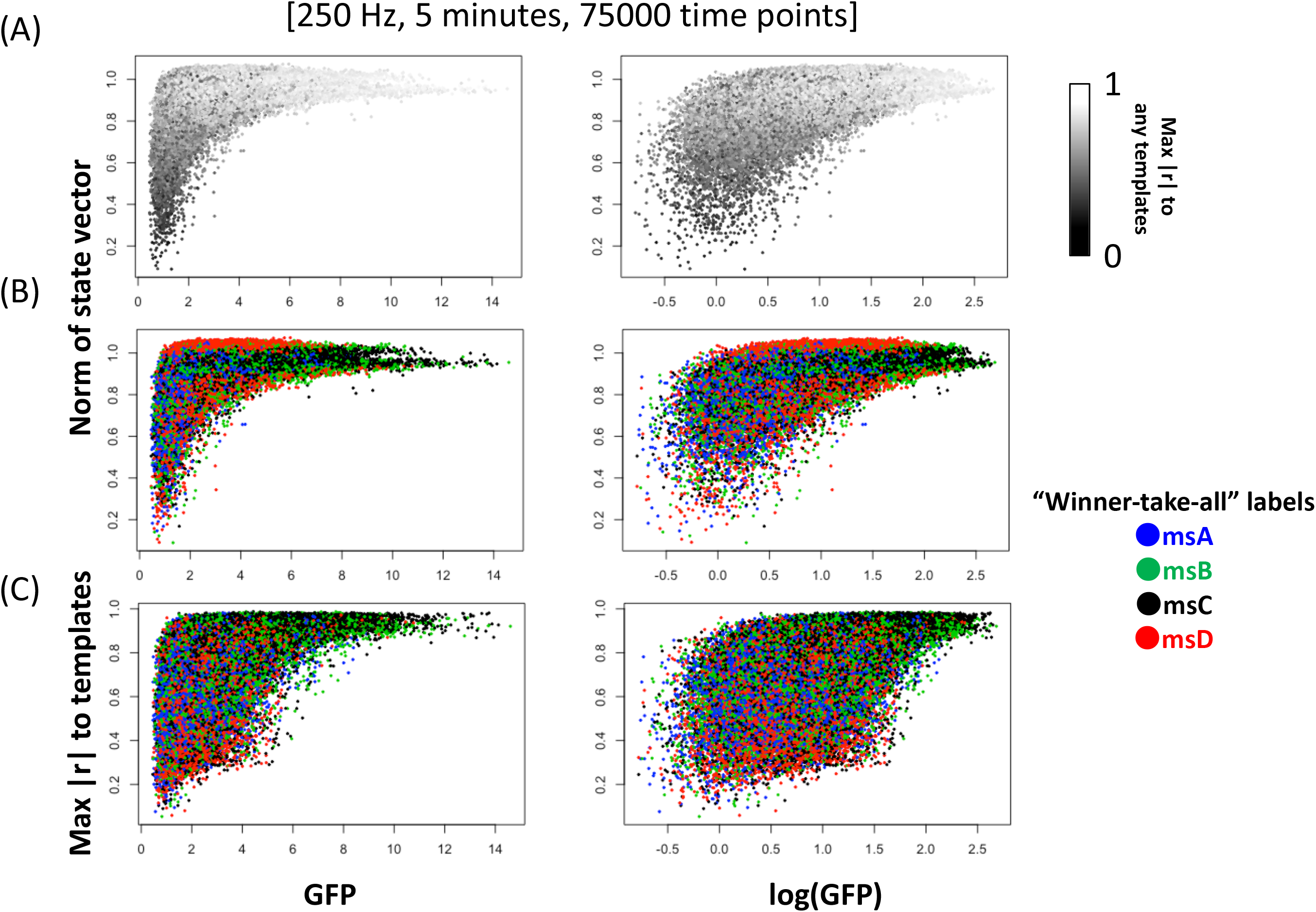
GFP, norms of state vector, and template-similarity.

### The globe as the common space

Thus far, we have suggested that the individual EEG state moves mainly on a spherical surface from one attractor to another (Fig. 3). If we consider that “sphericalness” is somehow related to the so-called “concentration on the sphere” in high dimensional space (Vershynin, 2018), the width of the surface might imply the latent dimensionality of EEG manifolds (the dimension is higher, the shell gets thinner). Theoretically, the grand-averaged EEG state (averaged over the participants) must be completely flat in voltage and located at the origin as an electrophysiologically impossible state. This could be analogous to the relationship between “average face” as a physiognomically rare phase (Grammer et al., 2002). However, when we are interested in the groupwise dynamics of the EEG state, the “hyperalignment” is required to match the individual spaces. Individual manifolds were depicted as spheres (Fig. 5). The number of templates (k problem in clustering) means how many parcels are expected for the surface (e.g., “four color theorem, (Appel et al., 1989) (Fig. 5AB for comparison between 4(8)- and 6(12)-template models with polarity) where the template state (“attractor”) should be located on the center in each region so that “winner-take-all” can be rephrased as “closer-take-all” (or, we selectively see template-near states only with high spatial correlations such as r > 0.8 in Fig. 5C). In addition, if necessary, we may create a polarity-ignored manifold from conventional absolute similarity (e.g., a spatial correlation matrix); figure 5D suggests that this is no longer a sphere. Furthermore, more templates or “landmarks” (in the current case) could be reproduced from the canonical templates as we created the polarized templates (Fig. 2B). The averaged state from the two (= 2 vectors) should be located approximately between their two positions on the spherical surface because they are all within a similarity space (Fig. 5E). These landmarks including canonical template-families can be further used to define the “standard space” where the following hyperalignment algorithm would run.

**Figure 5.**
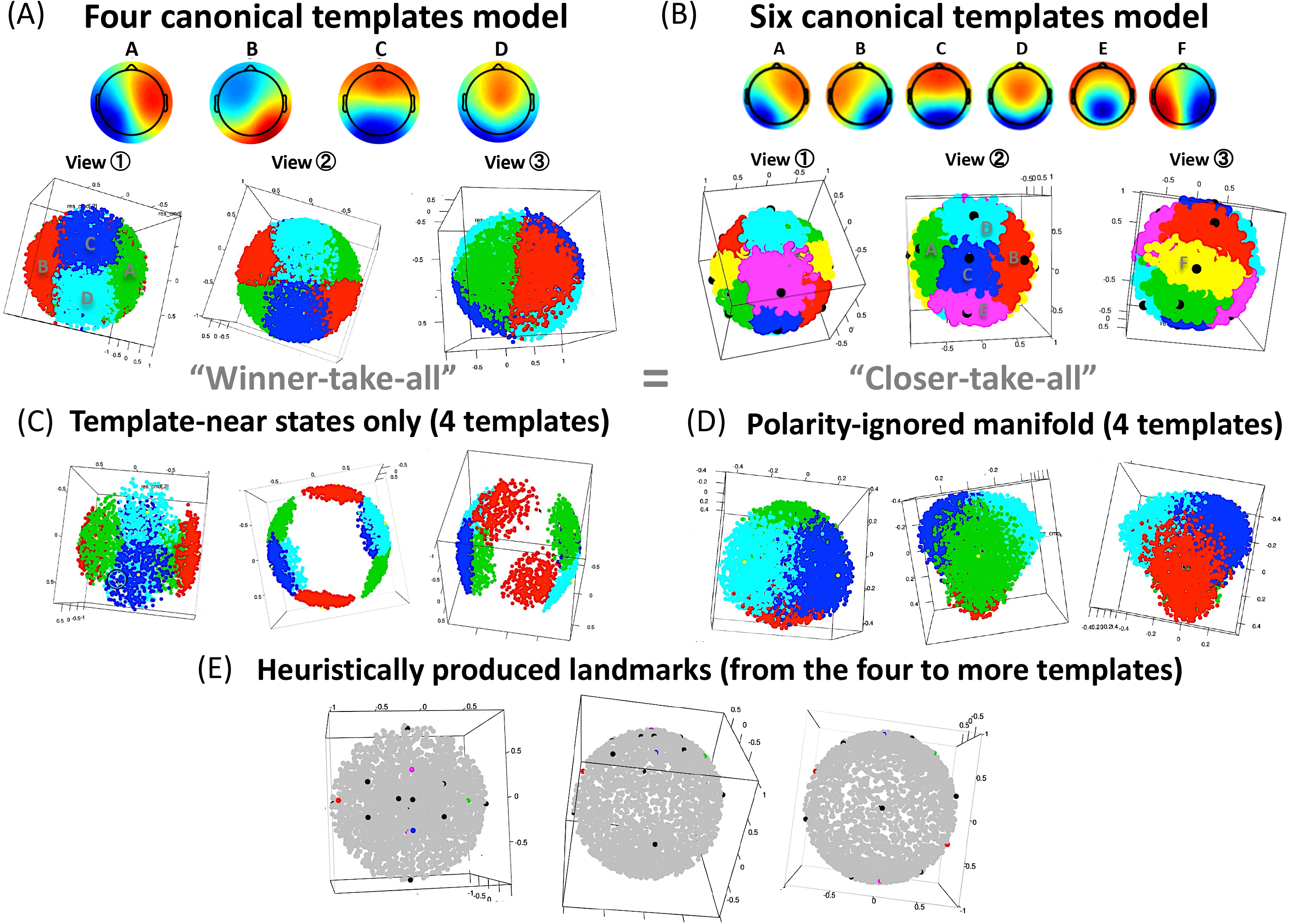
Variations of templates, manifolds, and landmarks.

Hyperaligning the registration of point groups over participants/sessions onto the same standard space is not difficult in the current case because the individual manifolds are all spheres with the same radius (each configuration is data-driven). Figure 6 shows our hyperalign procedure (“sequential MDS”) even for huge accumulated EEG states over participants/sessions. This was essential because applying MDS over grand-all states requires unrealistic computation time or immediately causes memory overflow, especially for electrophysiological EEG states (for example, a 100 Hz 5 min recording produces 30,000 states). First, we need “seeds” for the standard space. In a typical situation, some minutes of resting data for some participants are sufficient; for example, a 100 Hz recording for 1 min over five participants produces 30,000 states in total. Once the conventional MDS completes all these states, each point for each state shapes a sphere. Because not all points are required in the subsequent process (the more increases the accuracy of registration but requires more time), a sufficient number of points, such as 50 points, can be selected through grid sampling. Here, we can add templates (canonicals or canonical families, as mentioned above) to these and apply the MDS again over these states (e.g., 50 selected states and eight polarized templates). Although each position is no longer on the grid because the configuration is data-driven and the centroid is relocated to the origin (i.e., relative within the dataset), these can be used as our standard space (i.e., reapplying MDS to these does not change the configuration).

**Figure 6.**
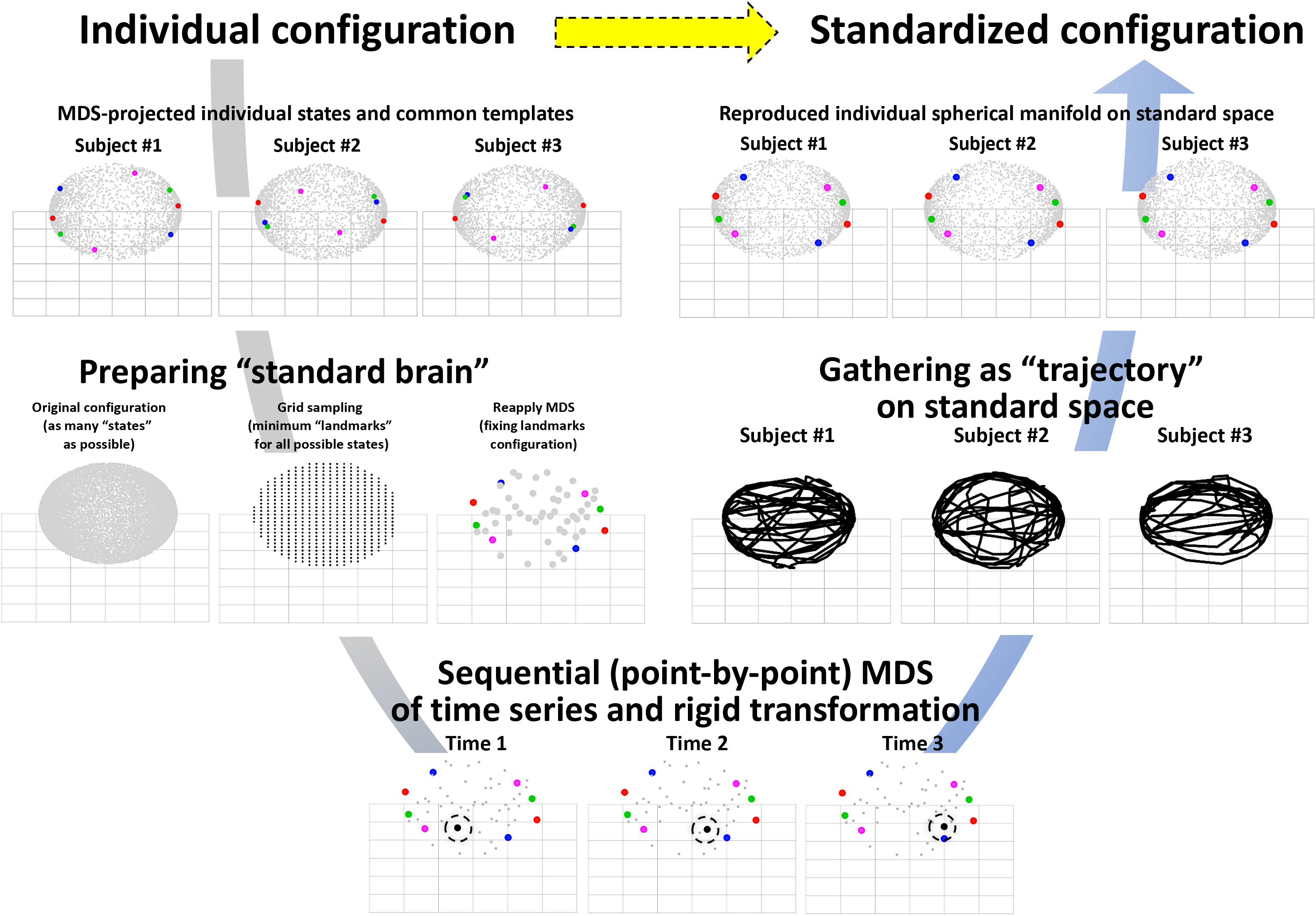
Hyper alignment over participants/sessions (sequential MDS)

We are now ready to apply point-by-point registration within the standard space (Fig. 6, lower). A new state was then added to the standard state (58 points). The MDS was applied to these 59 states and configured. If the number of standard points is large enough for one new point, the relative positions (i.e., distances) among the standard points do not change, although the configuration is slightly different from the original standard configuration (e.g., the centroid moves slightly). Therefore, a rigid transformation with reflection between the two configurations for standard states (original and temporally rearranged) can determine the position of the new state within the original standard space. Consequently, every new state can be located within the same space using this algorithm. This is a sequential process to allow as many states as possible to be located through sMDS (for example, for high-frequency sampled data) and is also useful for hyperalignment over participants. Another implementation is real-time neurofeedback to participants based on the determined position of the current state (xyz coordinates). Finally, we obtained individual but standardized manifolds for comparison (Fig. 6, upper right), where the templates were positioned at the same coordinates and the current state moved spatiotemporally around the templates in a continuous manner.

As a result of our sequential MDS algorithm, we can see and compare individual manifolds or trajectories even for an external large dataset where a different EEG cap with more channels was used (Babayan et al., 2019), and we were able to replicate it (Fig. 7A as view IZ for a typical participant). In addition, the state distribution is indicated as a heatmap from the same view (Fig. 7B) or a scatter plot on the sphere (Fig. 7C, when all norms are normalized to 1.0, or we can threshold shorter norms as noisy states). Furthermore, if we want to see the entire surface of the “globe” at a glance, various “map projection” methods are applicable with inevitable method-dependent biases (see also Supplementary Animation S1). Figure 7D shows the global field power (darker indicates higher GFP) for each state on the azimuthal equidistant projection (the distance and azimuth from the center are retained, so that the globe is represented as a perfect circle on the plane). Figure 7E represents four (eight) regions for each polarized canonical EEGms class (i.e., map projection of Fig. 7A), whereas Figure 7F is for Figure 7B. By analyzing a large dataset, we found that regardless of gender or age, individual manifolds always exhibit a sphere with a slight bias. Therefore, we detected mistakenly recorded EEG data by checking their individual spheres whose shapes were largely distorted (Fig. 7G-L). In addition, regarding preprocessing, we confirmed that only the band-pass filter is necessary (e.g., 2-20 Hz) for making spheres, although additional rigorous pipelines might help to modify noisy states that should be located near the origin of the space (Fig. 3B) as off-manifold “impossible” states (Sadtler et al., 2014). Figure 8 indicates another utility of spherical bias detection for our EEG-fMRI simultaneous recording data, where EEG signals were originally heavily biased by gradient and phase-encoding artifacts from the fMRI scans (Bullock et al., 2021; Chowdhury et al., 2019). Regardless of the single- or multi-band fMRI scans or phase-encoding directions (AP or PA), the largely biased manifolds were gradually restored toward a sphere through corrections and preprocessing stages (Fig. 8A, B, C, D).

**Figure 7.**
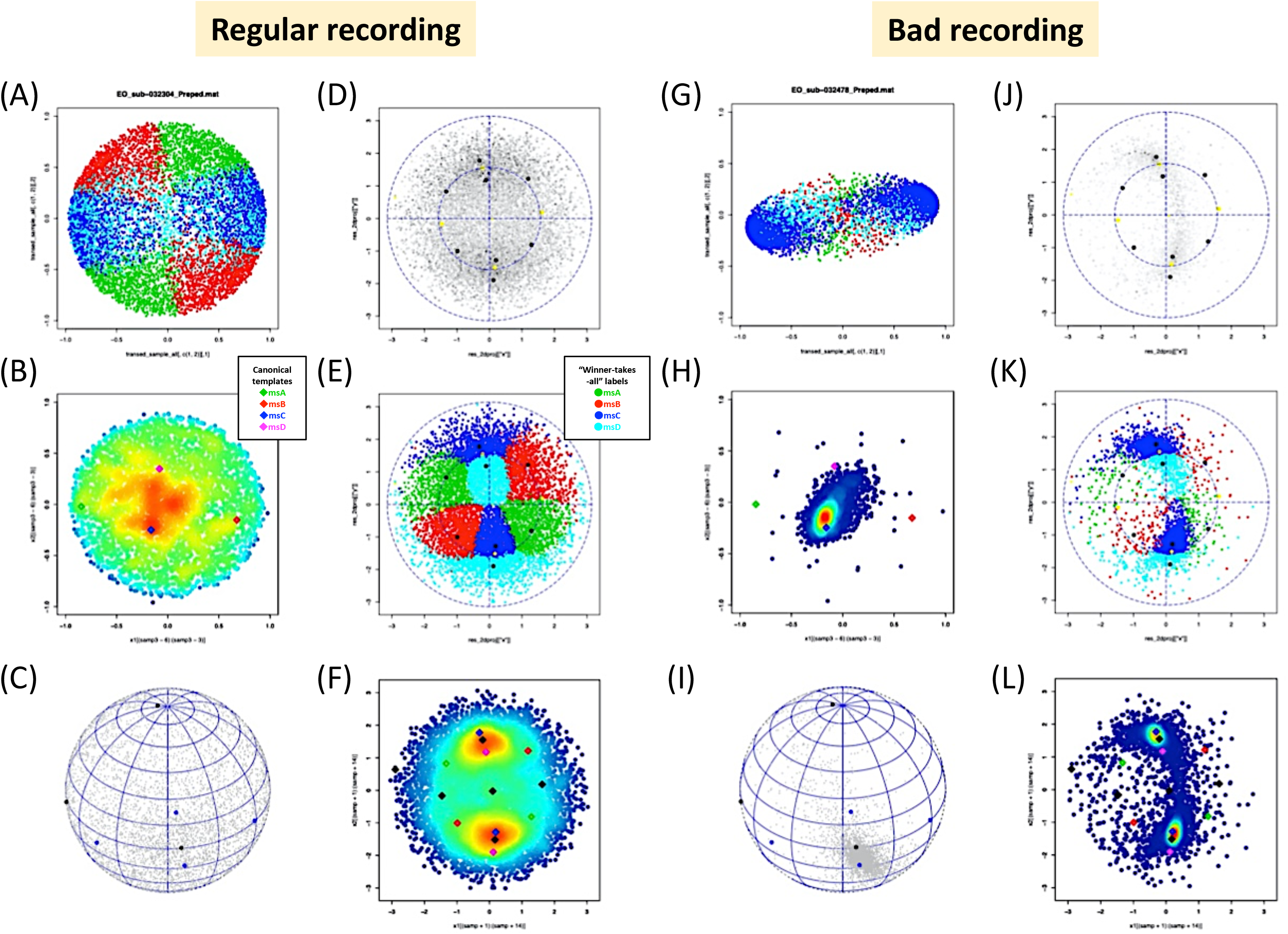
Robust individual spheres and useful spherical-bias detection

**Figure 8.**
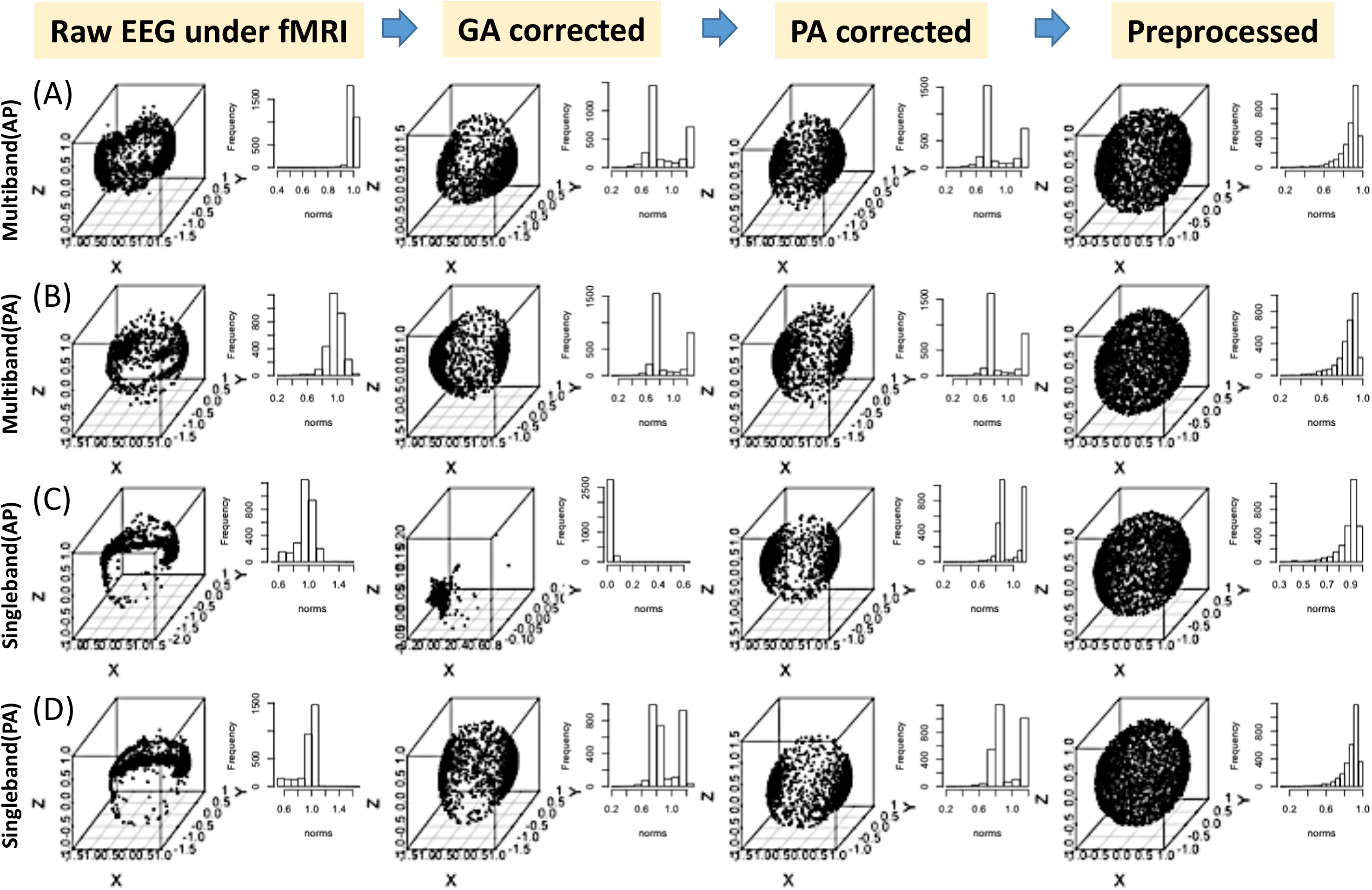
Spherical check of EEG signals under fMRI scans on each step.

### Discretization of continuous trajectories

A traditional EEGms analysis produces a transition probability matrix among canonical states or a state transition plot (e.g., a network plot), typically a 4 × 4 asymmetrical matrix and a directed graph with four nodes and 12 edges without self-recursive loops. These can then be updated (Fig. 9). First, we must polarize the states because the topographically positive and negative states are located on opposite sides of the sphere (Fig. 1). Second, we may increase the optimal number (k) of classes (Fig. 5B) because more classes represent more continuous trajectories (through a trade-off with coarse graining simplification). Finally, the directed graph can be depicted as pseudo 3D for the sphere because the positions of the nodes are determined through MDS, as mentioned above. Figure 9 illustrates the relationship among k, the transition matrix, and directed graphs. As expected, if k increases, the matrix becomes sparser (the transition route is fixed to some extent), but the graph represents a more spherical surface (e.g., k=1000), where we see an essential link between continuous and discrete pictures. The optimal k and the definition of the templates (“landmarks”) should be an important topic for future study, with going beyond some conventional statistical criterion. Figure 10 indicates age-related transition changes in the pseudosphere from a large external dataset (Babayan et al., 2019). Here, six landmarks (two for the north/south poles and four for the equator) were additionally reproduced from canonical 4 templates (Fig. 5E). The grand average of over 200 participants’ resting-state EEG suggests robust common trajectories (Fig. 10, upper left): essentially positive-negative transitions (i.e., between both topographical polarities) with specific A-B and C-D rotations. When the aged groups were subtracted from the grand average, C-D trajectories were characterized in our eyes: the younger as increased C-D rotations and the older as decreased C-D rotations (but increased trajectory on the equator).

**Figure 9.**
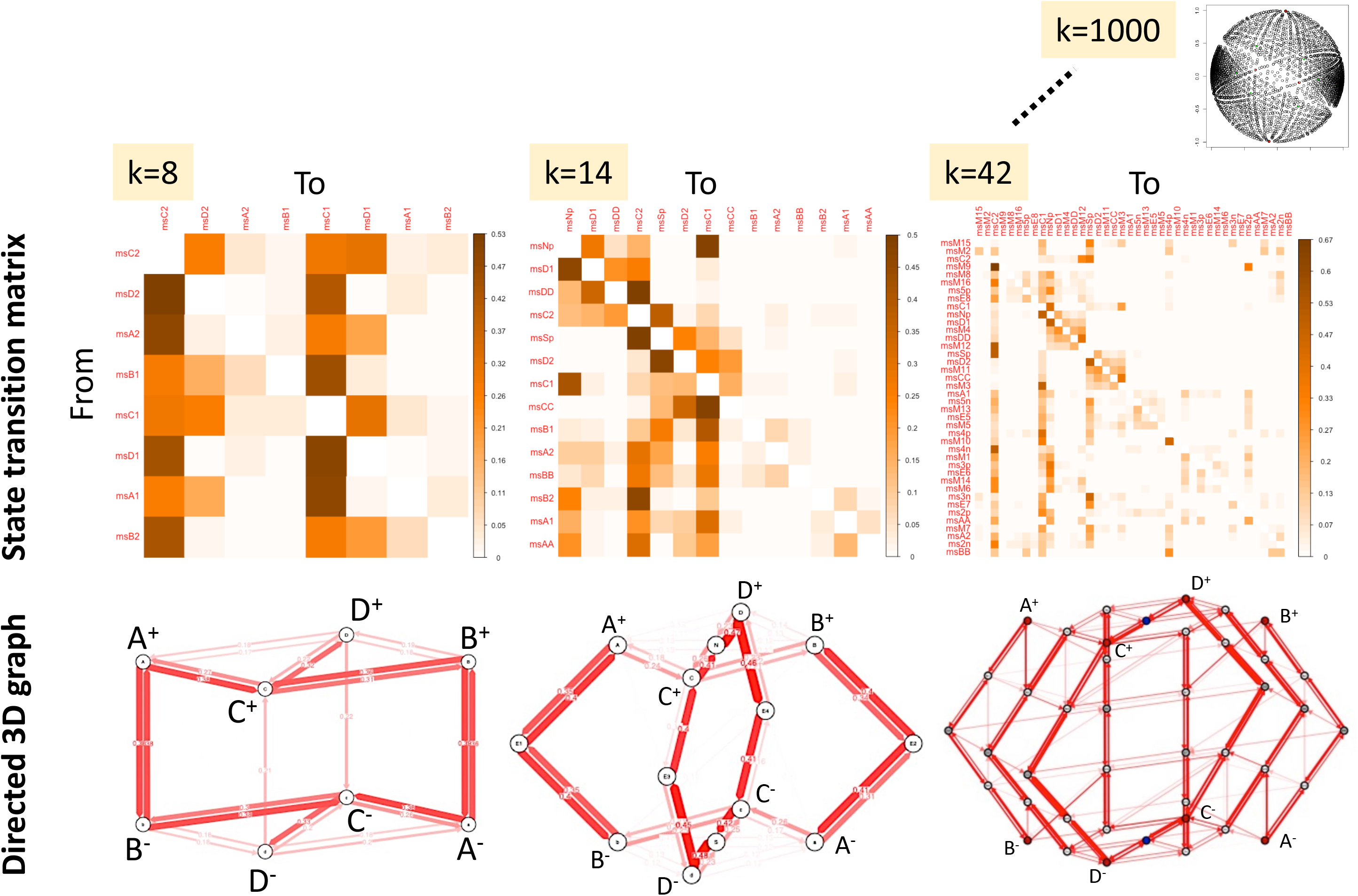
Exemplified state transition matrix and directed graphs (for pseudo sphere).

**Figure 10.**
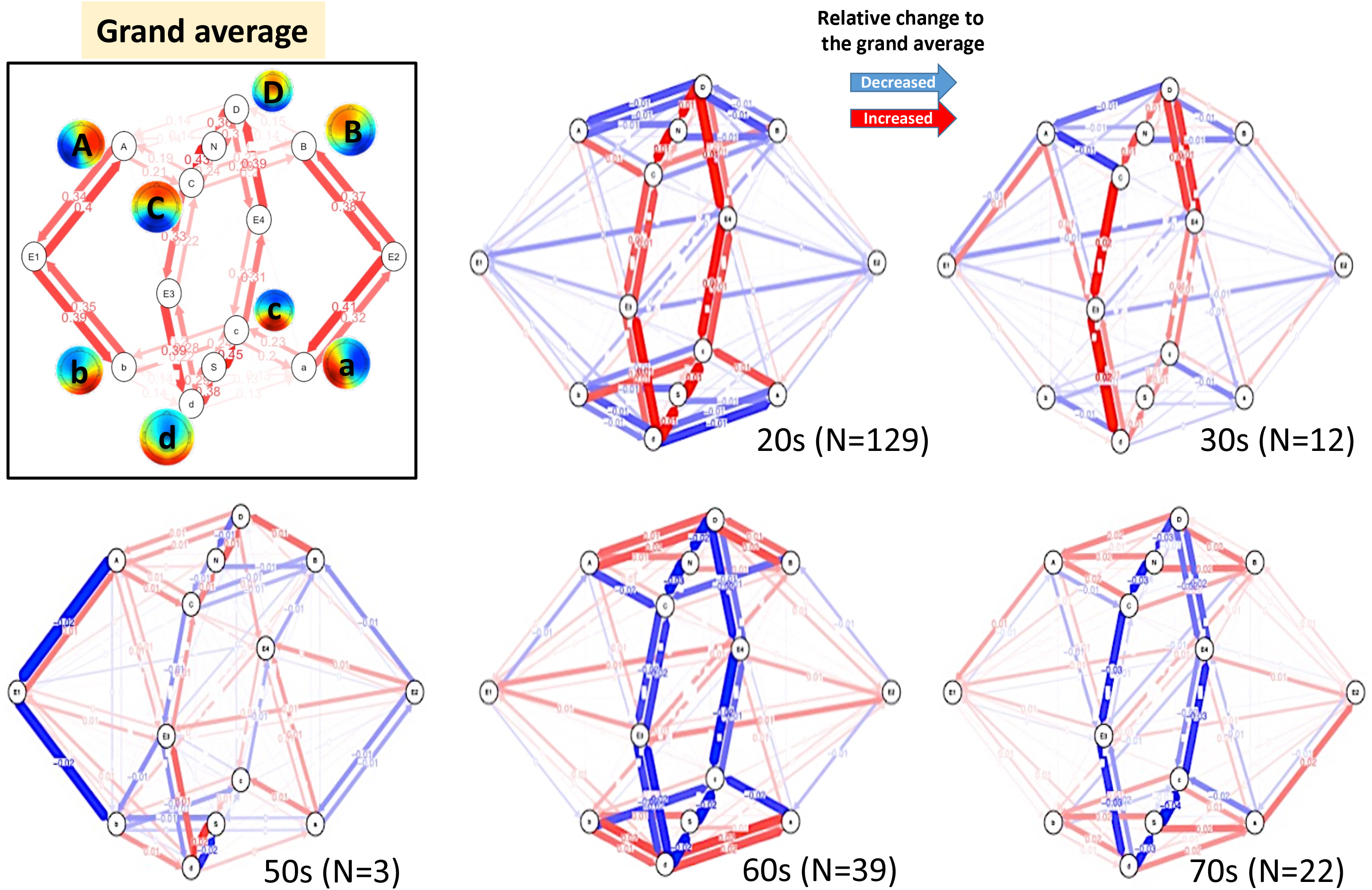
Age-related trajectorial changes on the pseudo-sphere.

### Modality independent procedure for the neural manifolds

Thus far, we have introduced a procedure for a standardized EEG manifold in which the current state is moving. This routine can be used as a modality-independent method for neural manifolds (Moon et al. 2019). Once the spatial correlation matrix was obtained (Fig. 2CD), MDS produced manifolds (Fig. 2E). Figure 11A represents MEG (Niso et al., 2016) manifold, where the state vector is, for example, 300D, whereas Figure 11B represents the fMRI manifold, where the state vector is approximately 900,000D, although both manifolds are embedded within the 3D similarity space without careful examination of its intrinsic dimensionality (see also Supplementary Animation S2). We observed ripples and spheres again in the minimally preprocessed data (e.g., bandpass filter) for both. In addition, if we can apply the same procedure to other neural manifolds, EEG microstate analysis may also be applicable for extracting attractor states (Ezaki et al., 2017). This may also be the case for MEG (Tait & Zhang, 2022) and fMRI. In the case of fMRI, the GFP peak dataset was also defined (Fig. 11C). After standard k-means clustering was performed, some templates (as cluster centroids) could be seen when k was set to 6 (Fig. 11D). Among these, a remarkable template should highly resemble a default mode network (DMN). However, our DMN template has some unique characteristics compared to those in the literature. First, our DMN is structure-oriented, although it is only an averaged low-resolution EPI image. Second, the contrast between activation suppression (red and blue) on the gray matter is much clearer. Third, “ polarized” DMNs (positive/negative) are extracted simultaneously (this is also true for other templates). Even after applying k-means over the spatially smoothed images (e.g., a 5 mm FWHM Gaussian filter), these characteristics might be maintained (Fig. 12 when k was 12). Since several whole-brain templates are available, the spatial similarity (Pearson’s correlation) with Smith ICA (independent component analysis (ICA) templates was depicted (Smith et al., 2009, 2013). Although the relationship is not one-by-one (Fig. 12A), our templates are congruent with the existing well-known network templates; however, ICA is presumably not for the definition of the momentary “states.” We minimally observed the default mode, executive, visual, and audio-motor networks (Fig. 12B for DMNs in the same plot definition).

**Figure 11.**
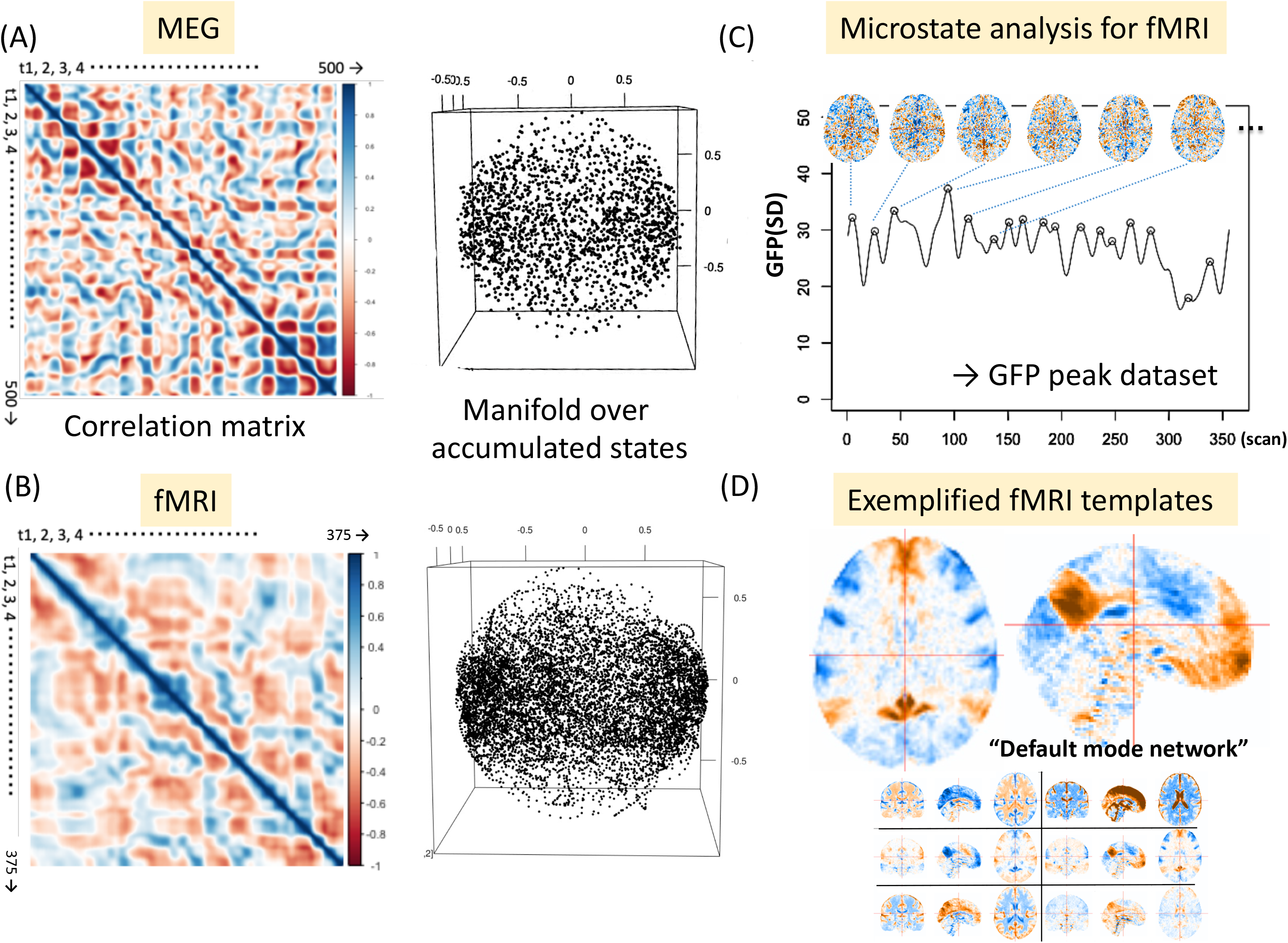
Modality-independent procedure.

**Figure 12.**
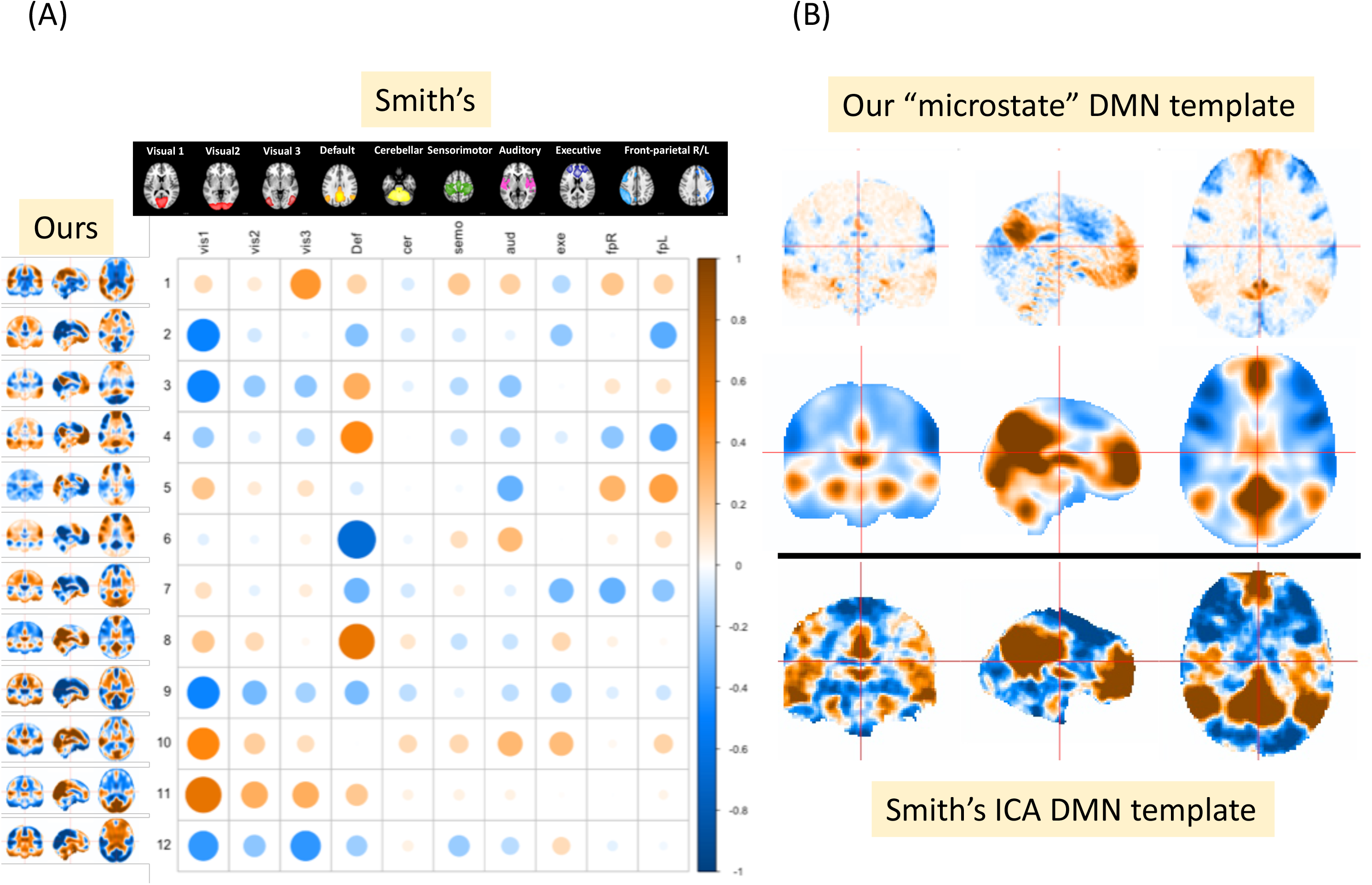
Spatially smoothed fMRI templates with Smith ICA templates

### Manifolds for phase-locked trajectories to be visualized

Finally, we suggest using this spherical neural manifold in future research. As our sphere was standardized, every state was located within the sphere for comparison. Therefore, we can compare simple trajectories rather than sensor-wise activities for experimental conditions, individual differences, and learning effects (Bradley et al., 2022; Iyer et al., 2022). Figure 13A shows the EEG manifold where some landmarks or “realms” are defined and condition-wise trajectories move around the regions (in this case, for the oddball task in Fig. 13A lowers) (see also Supplementary Animation S3). This is also applicable to the fMRI manifold (Fig. 13B for the n-back task) (see also Supplementary Animation S4). We now understand whole-brain neural dynamics both in a continuous and discrete manner (Fig. 2 and 9) as inseparable representations of the dynamic phase transition of the biological system.

**Figure 13.**
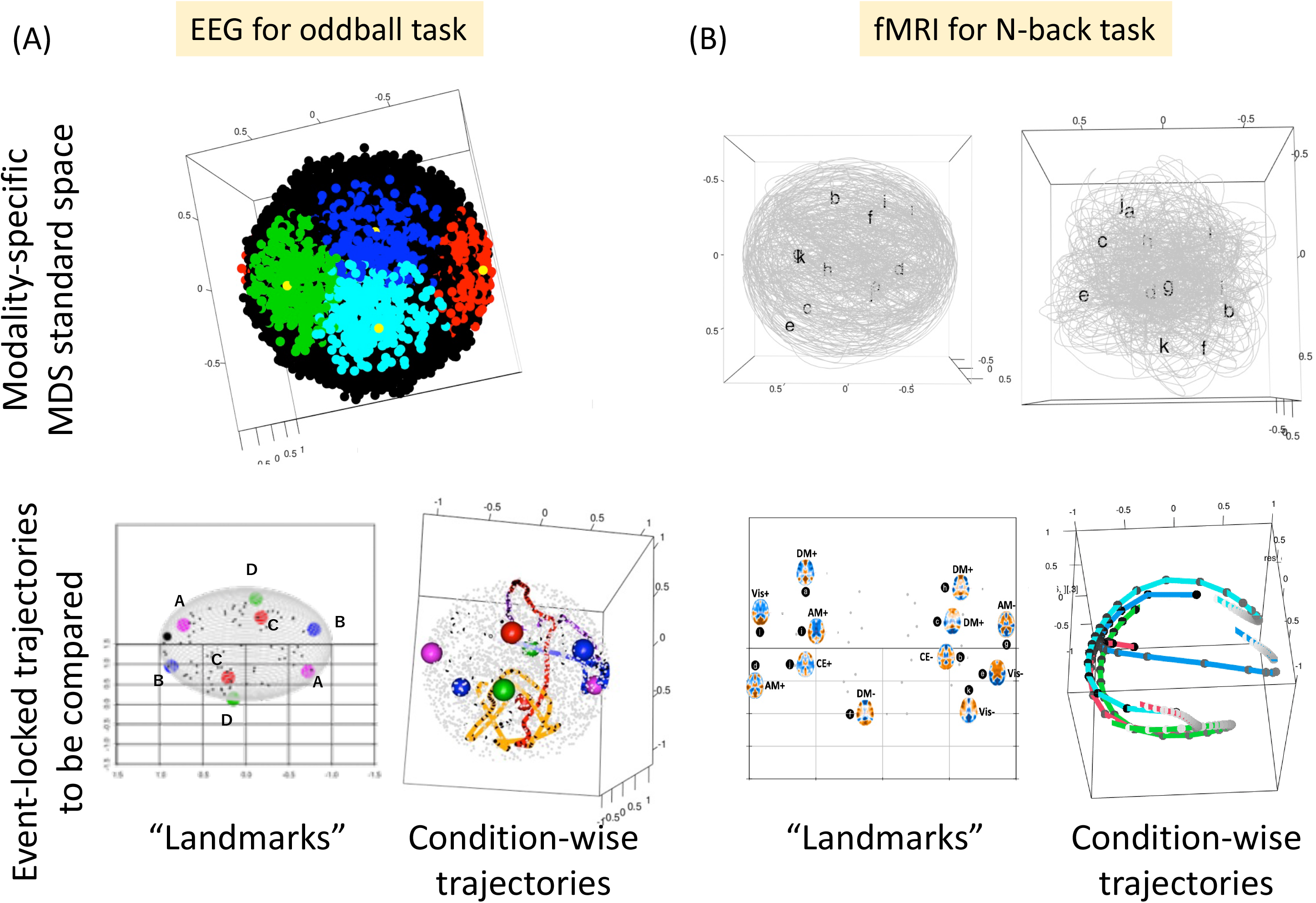
Event-locked (phase-locked) state dynamics on manifolds.

### Further specific insights into EEG/fMRI spherical manifold

In addition to its practical utility, the EEG/fMRI spherical manifold has provided significant insights. First, the visual representation of the distance matrix (see Fig. 2D), which clearly suggests its relevance to (nonlinear) neural dynamics, has potentially shared characteristics with fMRI/MEG (Fig. 11AB). Figure 14 depicts the intrinsic relationship between temporal and spatial autocorrelations in EEG. Further, when the distance matrix is depicted as unicolor (c.f., Fig. 2D), distinct checkerboard spatial patterns emerge in these matrices, with approximately 10 repetitions of on/off elements being remarkable (Fig. 14A). Where each state is defined as a simple voltage map, the repetitive on/off patterns indicate topographically polarized (reversely rotating) global state-transitions within a specific time window (Fig. 14B) (see also Supplementary Animation S5). Since these 10 repetitions might be linked with 10Hz alpha fluctuations (the most dominant for human brains), we investigated the effect of band-pass filters on the distance matrix. Hence, Figure 14C clearly depicts the presence of spatio-temporal homology, where channel-wise temporal filters determine the spatial dissimilarity (i.e., the fineness of checker patterns) (Supplementary Animation S6). These findings reveal the presence of various brain waves, including standing, traveling, and rotating waves, with their intrinsic relationship. For example, a broadband distance matrix could be obtained as a weighted summation of those from each narrow band, with the alpha matrix expected to be dominant. Since the distance matrix is calculated from voltage-free spatial patterns in this study (Supplementary Animation S7), in order to observe the behavior of waves (e.g., rotations), it is helpful to focus on the positive side (“wave crest”) after state-wise normalization (Supplementary Animation S8 left). Additionally, one can track the trajectory on the 2D unfolded state space, as depicted in Fig. 7DEF, where the rotation direction regularly changes to the opposite (Supplementary Animation S8 right). In addition, these phenomena could also be observable in the case of fMRI. For example, the global transitions between positive and negative DMN patterns become visible (Supplementary Animation S9 and S10).

**Figure 14.**
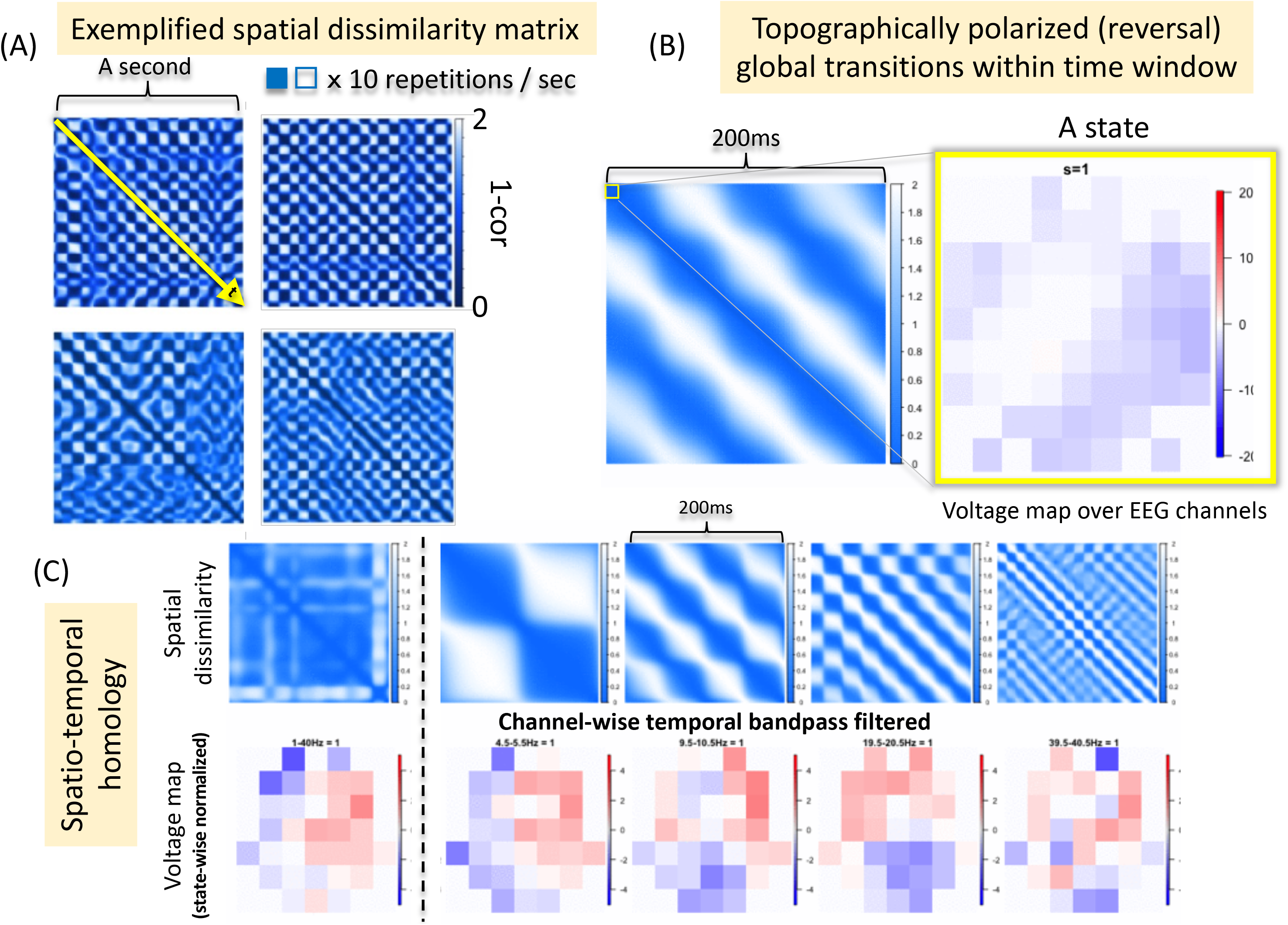
Intrinsic relationship between temporal and spatial auto-correlations.

Regarding fMRI manifolds, the fundamental characteristic or structure is shared with EEG manifolds, as we have partly suggested. However, we have identified a critical parameter for handling fMRI images to reproduce spherical surfaces: spatial smoothing. Figure 15 and 16 illustrate the effect of the spatial smoothing parameter (FWHM) after preprocessing (e.g., fMRIprep) on the subsequent analysis stage. In terms of achieving sphericalness with a radius of 1.0, more aggressive smoothing is found to be effective (Fig. 15), resulting in spatial representation that bears a close resemblance to EEG spatial representation achieved through channel interpolation (see Fig. 3B). Nevertheless, these aggressive smoothing comes at the cost of reduced spatial expressiveness. Therefore, the extracted DMN-like templates are visually valid only when applied to conservatively smoothed images (up to 10 mm, Fig. 16). The necessity of robust spatial smoothing might arise from residual individual anatomical differences even after preprocessing. Indeed, even with conservative smoothing, when examining spatial dissimilarity across participants, individual clusters still emerged on the diagonal (Fig. 15 lower). Therefore, to establish a standardized state space (spherical surface in this context), it is crucial to apply sufficient spatial smoothing (e.g., 10 mm), ensuring almost no individual differences while preserving spatial expressiveness (this holds true for surface only, Fig. 17). Furthermore, individual manifold and trajectory is also comparable between the pre- and post-spatial smoothing conditions (Supplementary Animation S11). Conclusively, exemplified resultant spherical surfaces are depicted for the whole brain (Fig. 18) or gray matter (Fig. 19), with extracted templates (labeled visually) representing the potential states across participants (for the trajectory, refer again to Supplementary Animation S10 right). This state space could encompass existing spatial representations, such as Smith’s ICA templates. The examination of potential topographical polarity, especially in fMRI studies conventionally (e.g., “height threshold”), has not been performed. However, our findings reveal that the reported spatial patterns (templates) are all configured within a hemisphere of the state space (Supplementary Animation S12 left). Further, if we incorporate the polarity-inverted templates, they are visibly located at antipodal coordinates to the origin (Supplementary Animation S12 right). Nonetheless, publicly available templates have been created differently, using different standard brains or preprocessing tools, leading to potential variations in the shape of the brain (Fig. 12B). Therefore, these pre-existing templates may not align with our surface but may be closer to the origin, unlike the case with data-driven templates (Fig. 18 and 19).

**Figure 15.**
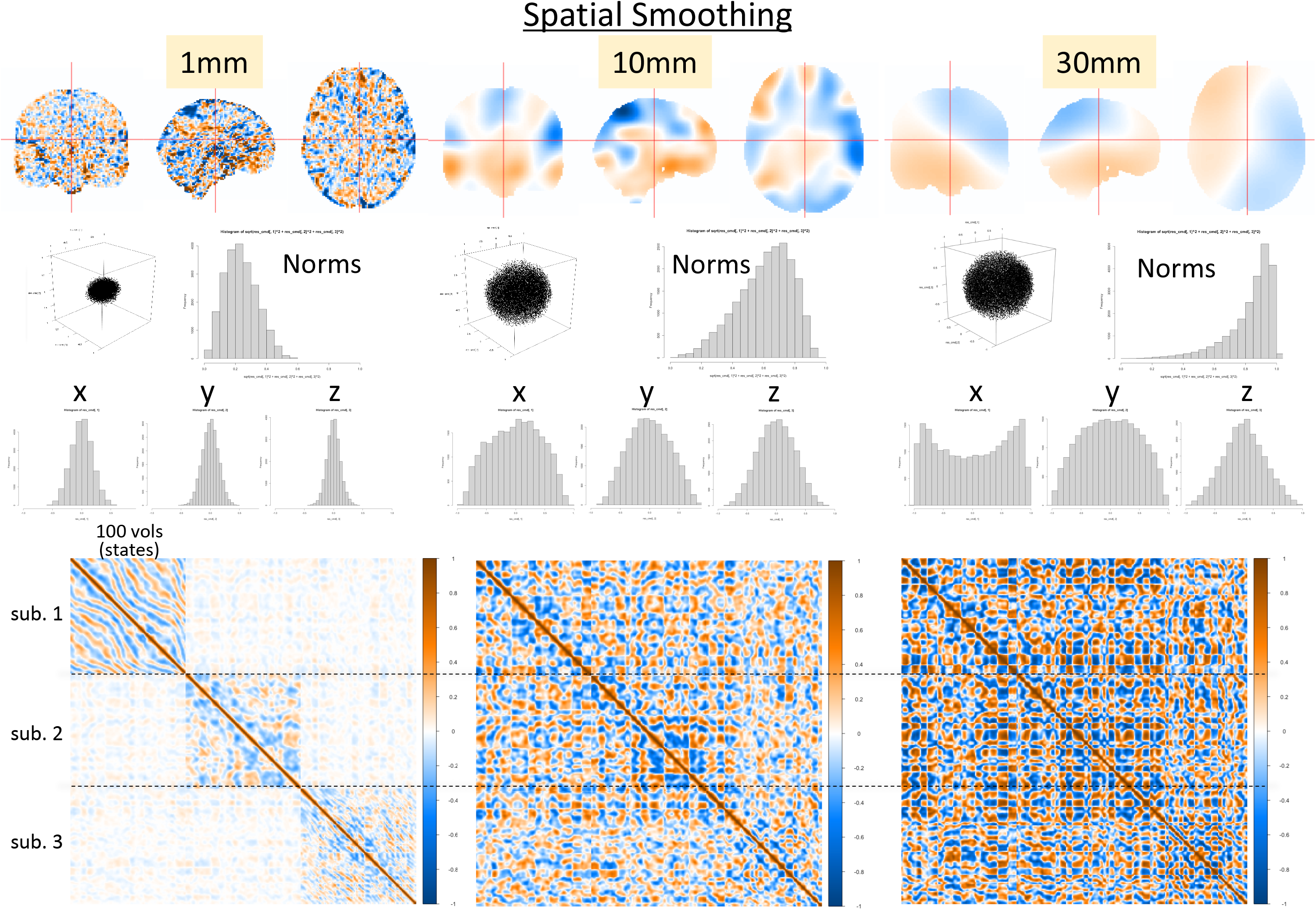
Robust spatial smoothing is necessary to normalize images and to reproduce spheres.

**Figure 16.**
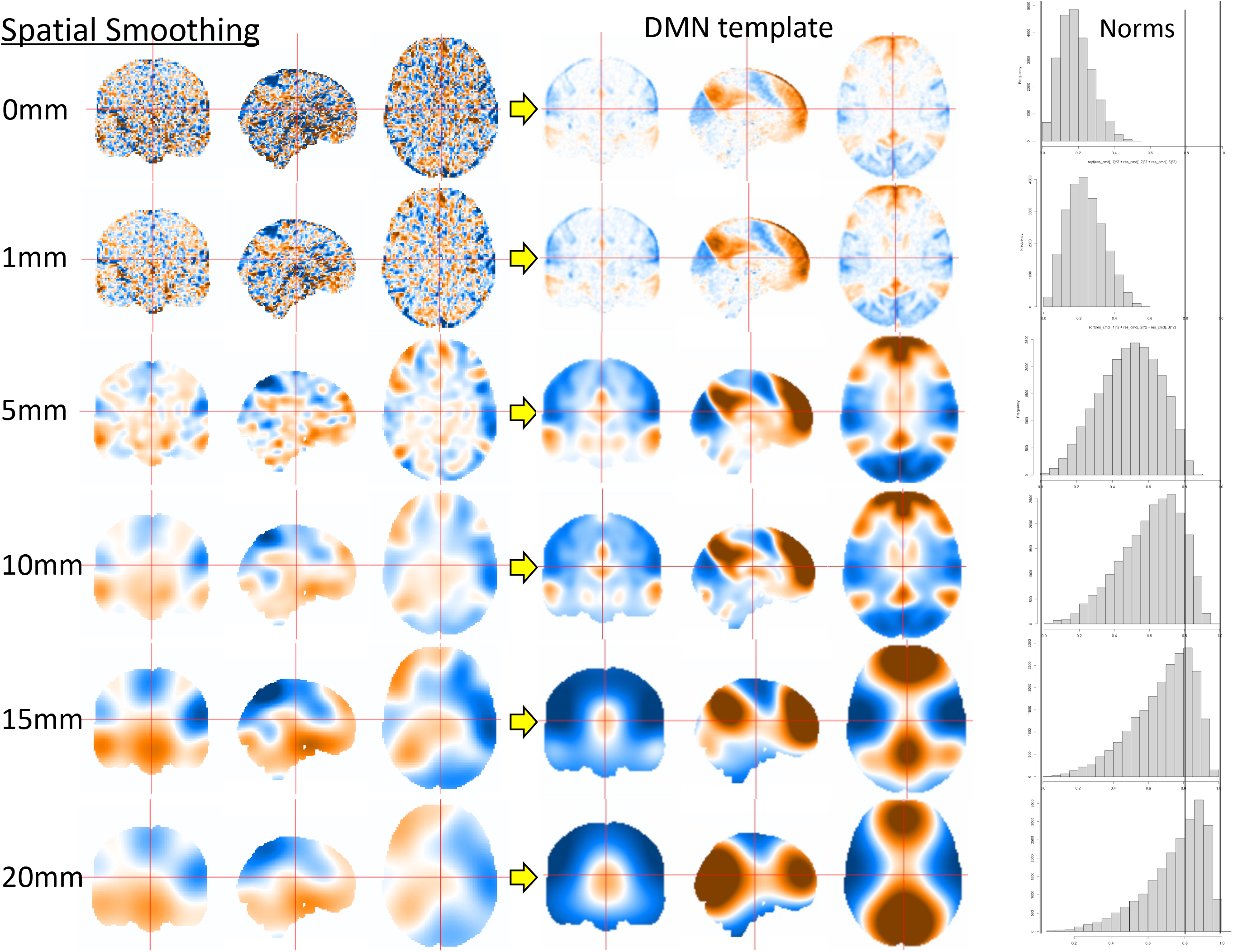
Trade-off between spatial smoothing (reducing regional specificity) and sphericalness.

**Figure 17.**
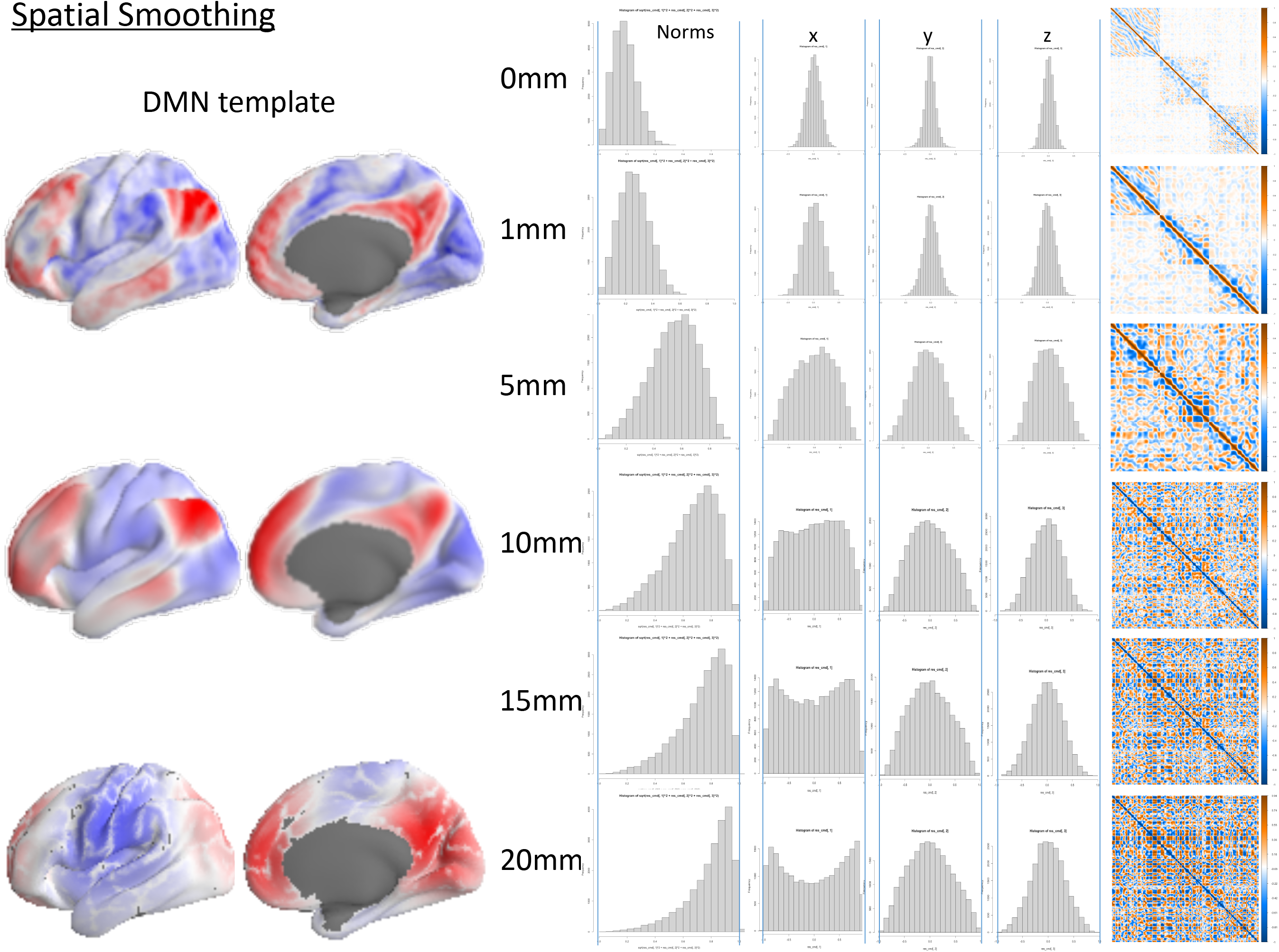
Surface (gray matter mask) exhibits the same trade-offs.

**Figure 18.**
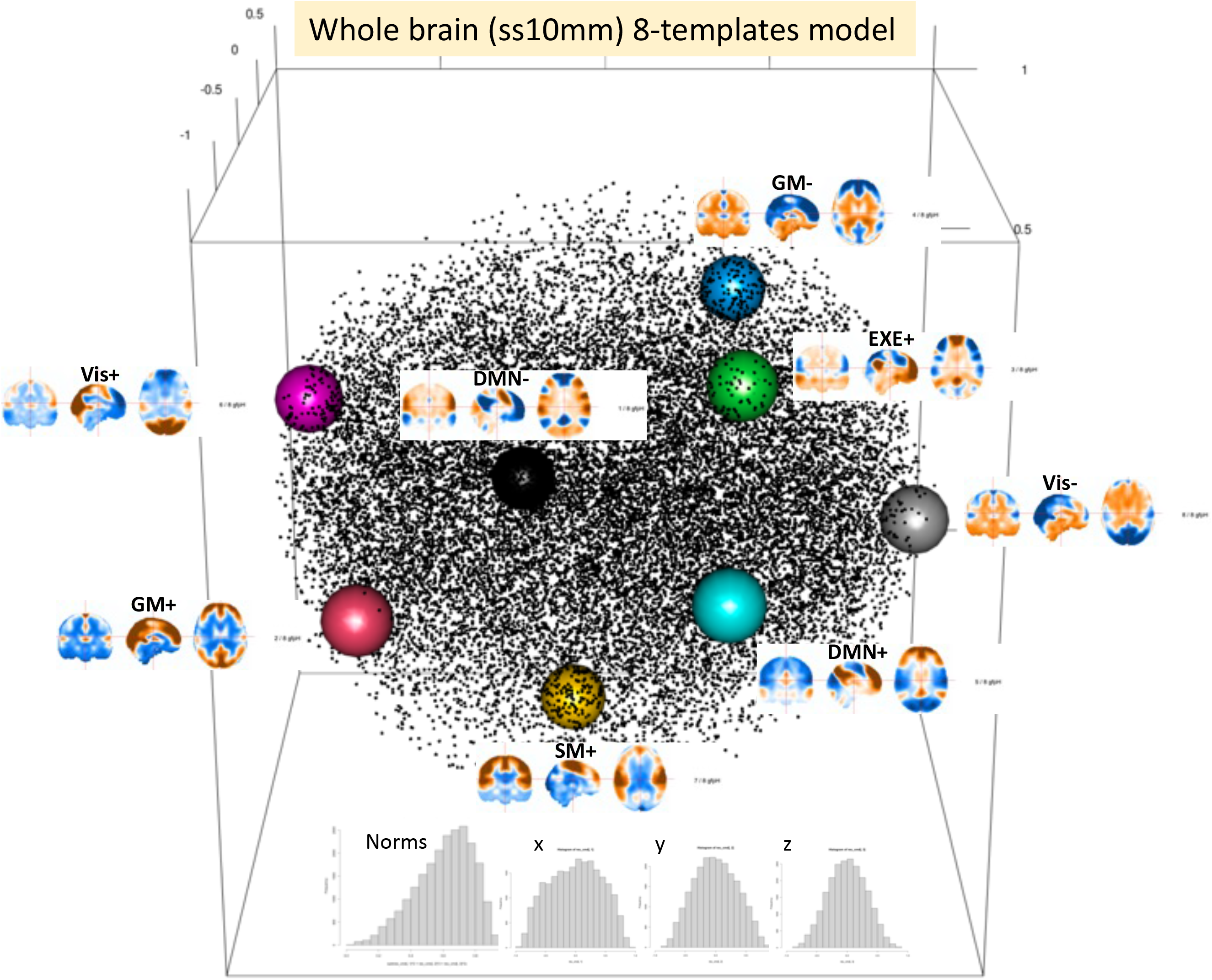
Whole-brain spherical state space with extracted 8 templates.

**Figure 19.**
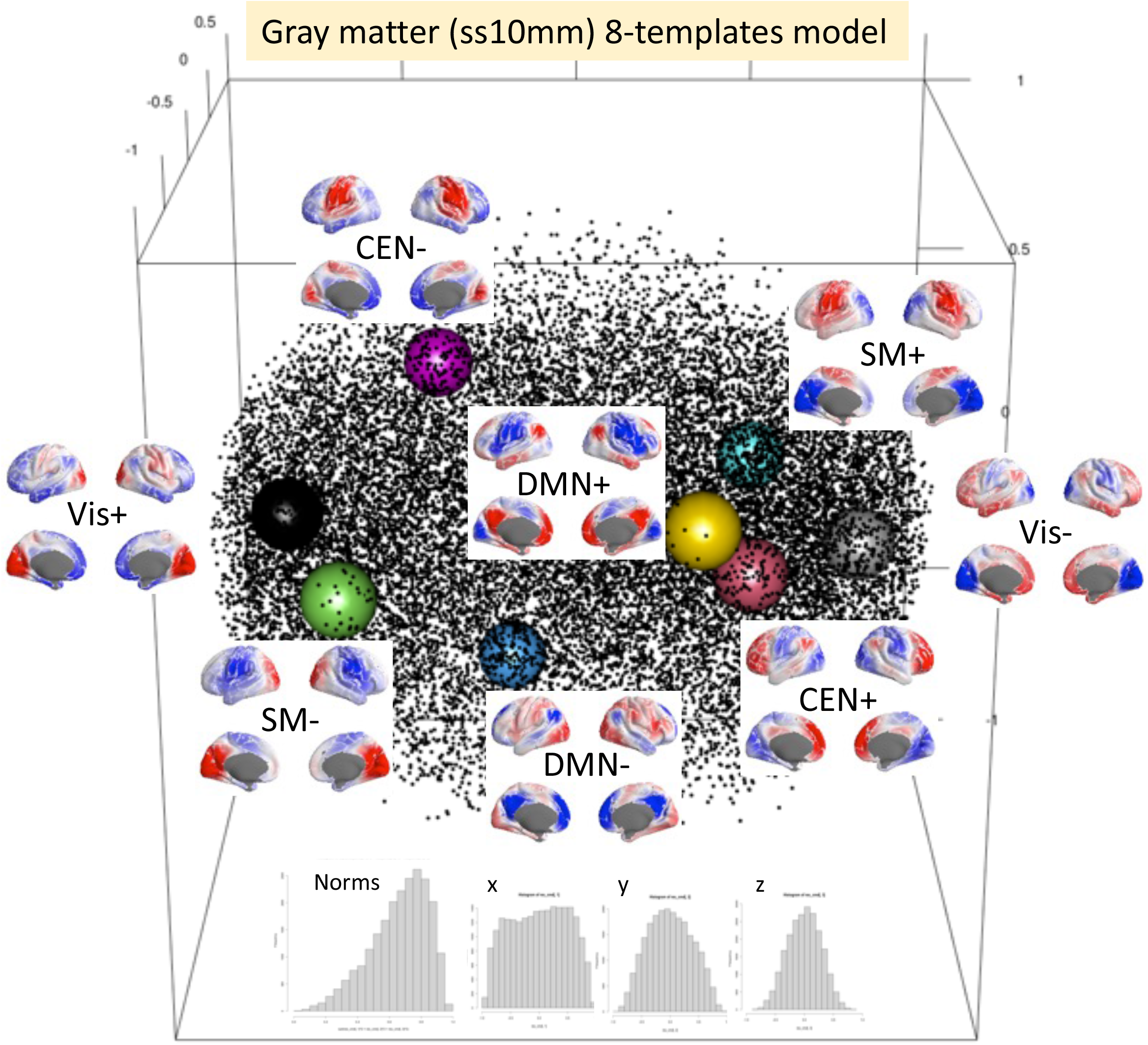
Gray-matter spherical state space with extracted 8 templates .

In conclusion, we propose a tentative depiction of state dynamics based on the spherical manifold, irrespective of neural modality. First, the local maxima of GFP provides a simple yet robust means of extracting attractor states. These states are traditionally assumed to best represent periods of momentary stability, reflecting the state of “activation” in the topography. In contrast, local minima are considered to be transient moments between states. For comparison, Figure 20 illustrates the distinction between the maxima and minima templates depending on the number of k in clustering, where the k2 maxima templates clearly suggest the global limits, akin to the north and south poles, exhibiting the highest GFP values globally (see Fig. 4BC again). Conversely, the k2 minima templates are located on the equator, forming a global basin. Our pilot analysis revealed that the timing of state changes driven by maxima templates (labeled using a winner-take-all approach) is closely aligned with the timing of GFP minima (this is a well-known phenomenon in EEGms analysis [back-fit results]). However, this is not observed in the results obtained using minima templates (e.g., k2 minima templates). Once we label each state, the transition matrix can be extended from two timepoints to include more timepoints (Fig. 21). By combining these matrices, the state transition can be chained, indicating a specific rotation with some non-directional hub nodes (i.e., multi-modal gradients, Supplementary Animation S13). Once the natural state space is defined, the position as input will generate the state as output (Supplementary Animation S14). Here, we can observe the correspondence between the rotation of the position within the state space (on the plane) and that of the peak voltage channel within the EEG topomap (Supplementary Animation S15). In addition, actual rotating state-dynamics (Supplementary Animation S8B) could be frequency-dependent as well as be age-dependent (Fig. 10 and Supplementary Animation S16). Exemplified individual spheres are plotted finally as interactive 3DCG to be manually compared (Supplementary Animation S17).

**Figure 20.**
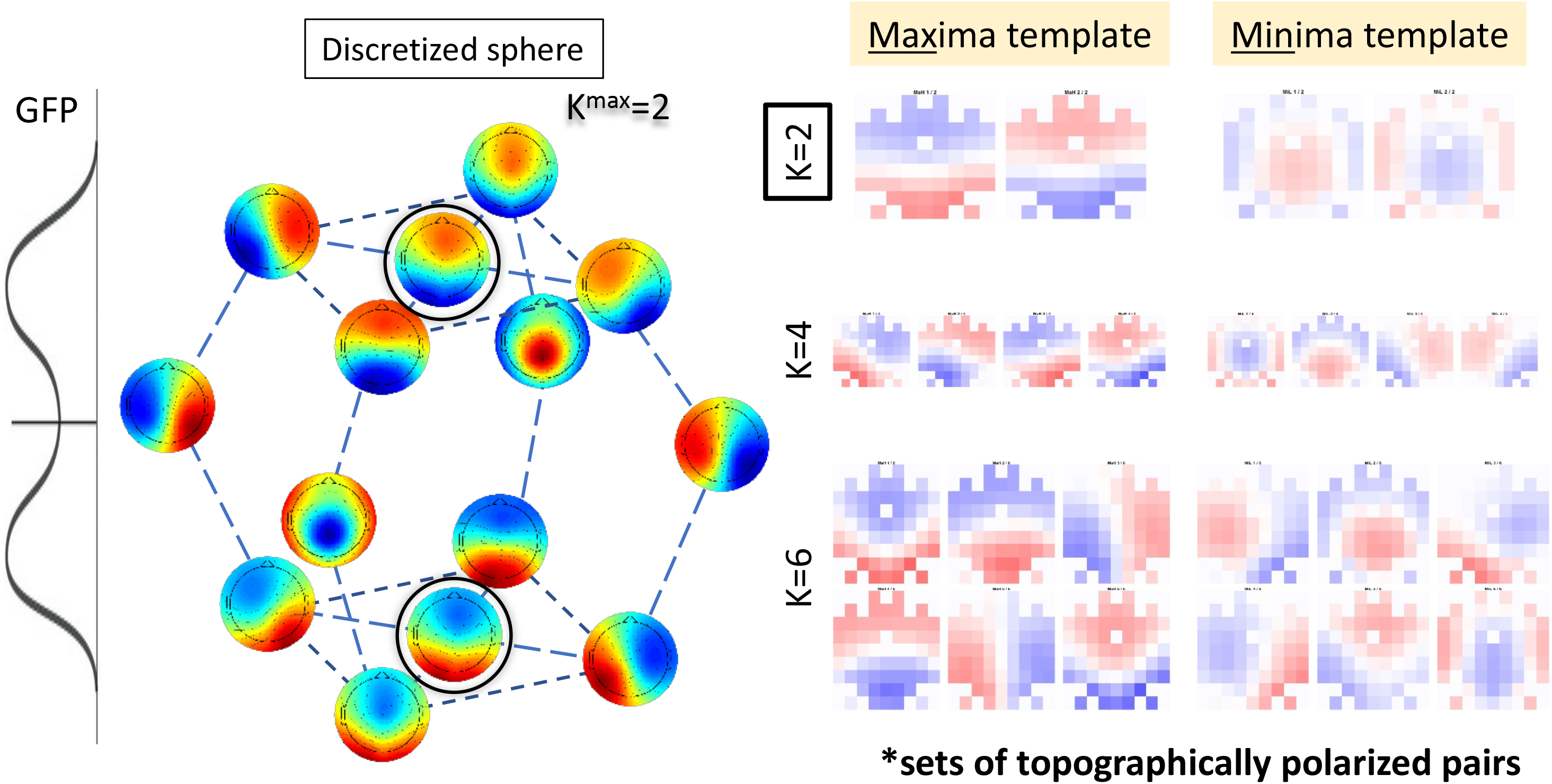
GFP maxima as local summits of mountain chains.

**Figure 21.**
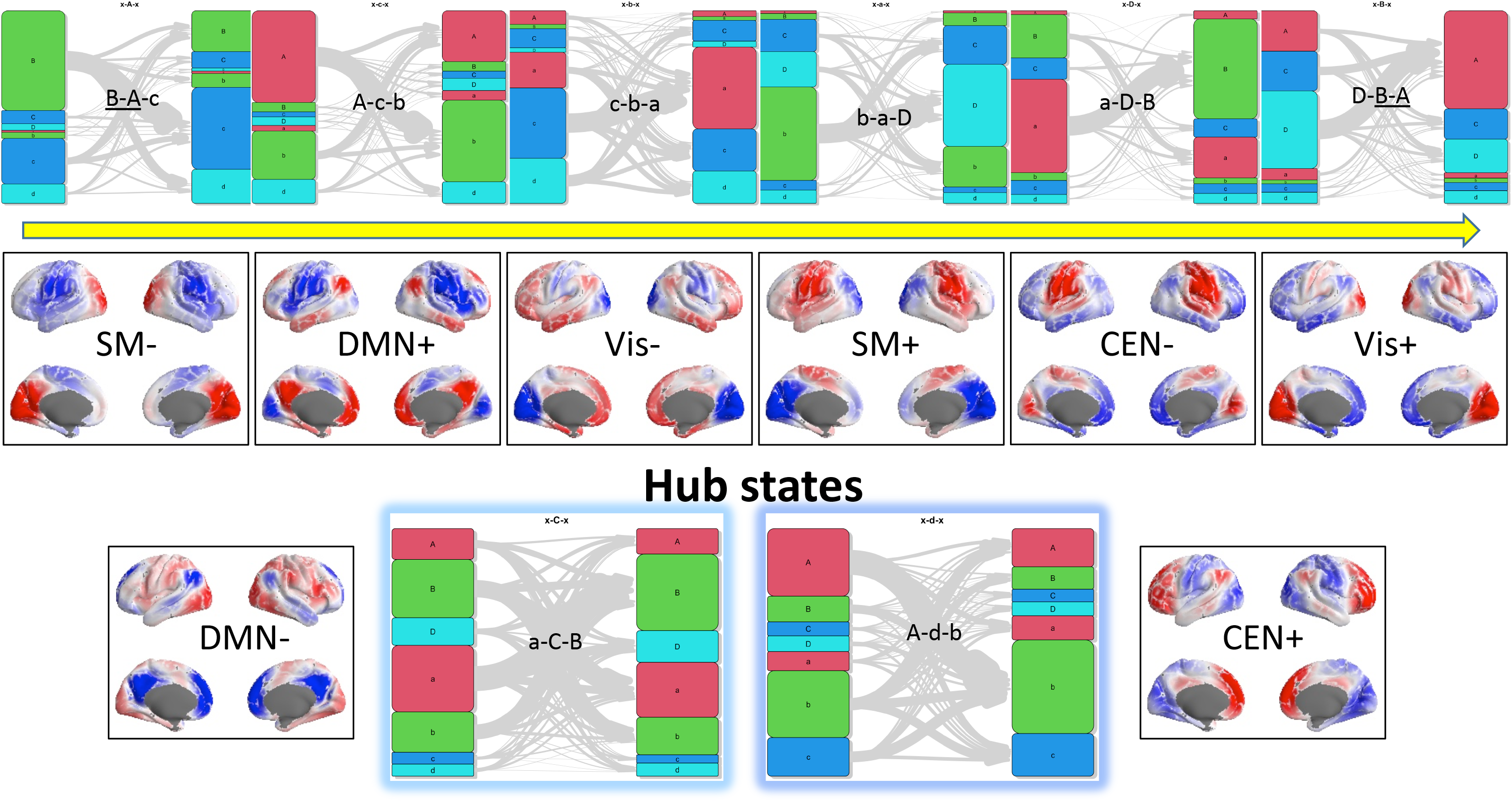
State-transitions could be chained with some hub nodes.

## Conclusions

The current study aimed to depict neural dynamics regardless of the measurement modality. Because neural dynamics are essentially continuous and thus infinite (i.e., a sphere), handling it is not easy without top-down mathematical approximate modeling. Our policy might be a bottom-up approach in which data-driven visualization is preferred. These results strongly imply continuous spatiotemporal dynamics on a spherical surface. Therefore, the problem is how to handle qualitative flow (i.e., global travelling). We suggest possible coarse-graining quantifications, including trajectories on the neural manifolds and the transition matrix between attractors. To validate and compare them further, we will demonstrate possible applications in our future work.

In this study, a chain of mountains is implicated as a neural landscape, where each mountain represents a distinct neural phase, much like how each real mountain is uniquely named (while only some valleys have names). Climbers instinctively aim to ascend the local summits within these mountain chains. We now recognize that all mountains are interconnected globally, forming a unified entity. The “global” state dynamics could be illustrated through the lens of two songs: “Climb Ev’ry Mountain (in English lyrics)” representing a first-person active perspective and “Carrying You (in Japanese lyrics)” representing a third-person passive perspective, within the framework of an agent-world dynamical system.

## Supplementary Material

The supplementary animations as s single mp4 video are available online: https://www.dropbox.com/scl/fi/994ajntamkhpp9agi7dmf/supplementary-animations.mp4?rlkey=d10ho0yqjh8ekq3usedmy05yk&dl=0

## Acknowledgement

This study was supported by the Innovative Science and Technology Initiative for Security (Grant Number JPJ004596), ATLA, Japan.

## References

Appel, K, Appel KI, Haken W. Every planar map is four colorable. American Mathematical Society; 1989

Asai T, Hamamoto T, Kashihara S, Imamizu H. Real-time detection and feedback of canonical electroencephalogram microstates: Validating a neurofeedback system as a function of delay. Front Syst Neurosci. 16, 786200 (2022). https://doi.org/10.3389/fnsys.2022.786200

Babayan A, et al. A mind-brain-body dataset of MRI, EEG, cognition, emotion, and peripheral physiology in young and old adults. Sci Data. 6, 180308 (2019). https://doi.org/10.1038/sdata.2018.308

Bagdasarov A, Roberts K, Bréchet L, Brunet D, Michel CM, Gaffrey MS. Spatiotemporal dynamics of EEG microstates in four- to eight-year-old children: Age- and sex-related effects. Dev Cogn Neurosci. 57, 101134 (2022). https://doi.org/10.1016/j.dcn.2022.101134

Barack DL, Krakauer JW. Two views on the cognitive brain. Nat Rev Neurosci. 22, 359–71 (2021). https://doi.org/10.1038/s41583-021-00448-6

Bradley C, Nydam AS, Dux PE, Mattingley JB. State-dependent effects of neural stimulation on brain function and cognition. Nat Rev Neurosci. 23, 459–75 (2022). https://doi.org/10.1038/s41583-022-00598-1

Bullock M, Jackson GD, Abbott DF. Artifact reduction in simultaneous EEG-fMRI: A systematic review of methods and contemporary usage. Front Neurol. 12, 622719 (2021). https://doi.org/10.3389/fneur.2021.622719

Chowdhury MEH, Khandakar A, Mullinger KJ, Al-Emadi N, Bowtell R. Simultaneous EEG-fMRI: Evaluating the effect of the EEG cap-cabling configuration on the gradient artifact. Front Neurosci. 13, 690 (2019). https://doi.org/10.3389/fnins.2019.00690

da Cruz JR, et al. EEG microstates are a candidate endophenotype for schizophrenia. Nat Commun. 11, 3089 (2020). https://doi.org/10.1038/s41467-020-16914-1

de Bock R, Mackintosh AJ, Maier F, Borgwardt S, Riecher-Rössler A, Andreou C. EEG microstates as biomarker for psychosis in ultra-high-risk patients. Transl Psychiatry. 10, 300 (2020). https://doi.org/10.1038/s41398-020-00963-7

Ezaki T, Watanabe T, Ohzeki M, Masuda N. Energy landscape analysis of neuroimaging data. Philos Trans A Math Phys Eng Sci. 375 (2017). https://doi.org/10.1098/rsta.2016.0287

Férat V, et al. Electroencephalographic microstates as novel functional biomarkers for adult attention-deficit/hyperactivity disorder. Biol Psychiatry Cogn Neurosci Neuroimaging. 7, 814–23 (2022). https://doi.org/10.1016/j.bpsc.2021.11.006

Friston K, Moran RJ, Nagai Y, Taniguchi T, Gomi H, Tenenbaum J. World model learning and inference. Neural Netw. 144, 573–90 (2021). https://doi.org/10.1016/j.neunet.2021.09.011

Grammer K, Fink B, Juette A, Ronzal G, Thornhill R. Female faces and bodies: N-dimensional feature space and attractiveness. Facial Attractiveness Evol Cogn Soc Perspect. 311, 91–125 (2002).

Iyer KK, et al. Focal neural perturbations reshape low-dimensional trajectories of brain activity supporting cognitive performance. Nat Commun. 13, 4 (2022). https://doi.org/10.1038/s41467-021-26978-2

Jazayeri M, Afraz A. Navigating the neural space in search of the neural code. Neuron. 93, 1003–14 (2017). https://doi.org/10.1016/j.neuron.2017.02.019

Koenig T, Brandeis D. Inappropriate assumptions about EEG state changes and their impact on the quantification of EEG state dynamics. NeuroImage. 125, 1104–6 (2016). https://doi.org/10.1016/j.neuroimage.2015.06.035

Kriegeskorte N, Wei XX. Neural tuning and representational geometry. Nat Rev Neurosci. 22, 703–18 (2021). https://doi.org/10.1038/s41583-021-00502-3

Lehmann D. Multichannel topography of human alpha EEG fields. Electroencephalogr Clin Neurophysiol. 31, 439–49 (1971). https://doi.org/10.1016/0013-4694(71)90165-9

Mathys C, Daunizeau J, Friston KJ, Stephan KE. A bayesian foundation for individual learning under uncertainty. Front Hum Neurosci. 5, 39 (2011). https://doi.org/10.3389/fnhum.2011.00039

Michel CM, Koenig T. EEG microstates as a tool for studying the temporal dynamics of whole-brain neuronal networks: A review. NeuroImage. 180, 577–93 (2018). https://doi.org/10.1016/j.neuroimage.2017.11.062

Milz P, Pascual-Marqui RD, Achermann P, Kochi K, Faber PL. The EEG microstate topography is predominantly determined by intracortical sources in the alpha band. NeuroImage. 162, 353–61 (2017). https://doi.org/10.1016/j.neuroimage.2017.08.058

Mishra A, Englitz B, Cohen MX. EEG microstates as a continuous phenomenon. NeuroImage. 208, 116454 (2020). https://doi.org/10.1016/j.neuroimage.2019.116454

Moon KR, et al. Visualizing structure and transitions in high-dimensional biological data. Nat Biotechnol. 37, 1482–92 (2019). https://doi.org/10.1038/s41587-019-0336-3

Nieh EH, et al. Geometry of abstract learned knowledge in the hippocampus. Nature. 595, 80–4 (2021). https://doi.org/10.1038/s41586-021-03652-7

Niso G, et al. OMEGA: The open MEG archive. NeuroImage. 124, 1182–7 (2016). https://doi.org/10.1016/j.neuroimage.2015.04.028

Pascual-Marqui RD, Michel CM, Lehmann D. Segmentation of brain electrical activity into microstates: Model estimation and validation. IEEE Trans Biomed Eng. 42, 658–65 (1995). https://doi.org/10.1109/10.391164

Perrottelli A, Giordano GM, Brando F, Giuliani L, Mucci A. EEG-based measures in at-risk mental state and early stages of schizophrenia: A systematic review. Front Psychiatry. 12, 653642 (2021). https://doi.org/10.3389/fpsyt.2021.653642

Ros T, J Baars B, Lanius RA, Vuilleumier P. Tuning pathological brain oscillations with neurofeedback: A systems neuroscience framework. Front Hum Neurosci. 8, 1008 (2014). https://doi.org/10.3389/fnhum.2014.01008

Sadtler PT, et al. Neural constraints on learning. Nature. 512, 423–6 (2014). https://doi.org/10.1038/nature13665

Shaw SB, Dhindsa K, Reilly JP, Becker S. Capturing the forest but missing the trees: Microstates inadequate for characterizing shorter-scale EEG dynamics. Neural Comput. 31, 2177–211 (2019). https://doi.org/10.1162/neco_a_01229

Smith SM, et al. Correspondence of the brain’s functional architecture during activation and rest. Proc Natl Acad Sci U S A. 106, 13040–5 (2009). https://doi.org/10.1073/pnas.0905267106

Smith SM, et al. Functional connectomics from resting-state fMRI. Trends Cogn Sci. 17, 666–82 (2013). https://doi.org/10.1016/j.tics.2013.09.016

Tait L, Zhang J. MEG cortical microstates: Spatiotemporal characteristics, dynamic functional connectivity and stimulus-evoked responses. NeuroImage. 251, 119006 (2022). https://doi.org/10.1016/j.neuroimage.2022.119006

Vershynin R. High-dimensional probability: An introduction with applications in data science. Cambridge University Press 2018

von Wegner F, Knaut P, Laufs H. EEG microstate sequences from different clustering algorithms are information-theoretically invariant. In: Front Comput Neurosci. 12, 70 (2018). https://doi.org/10.3389/fncom.2018.00070

Watanabe T, Masuda N, Megumi F, Kanai R, Rees G. Energy landscape and dynamics of brain activity during human bistable perception. Nat Commun. 5, 4765 (2014). https://doi.org/10.1038/ncomms5765

Zanesco AP. EEG electric field topography is stable during moments of high field strength. Brain Topogr. 33, 450–60 (2020). https://doi.org/10.1007/s10548-020-00780-7

Zanesco AP, King BG, Skwara AC, Saron CD. Within and between-person correlates of the temporal dynamics of resting EEG microstates. NeuroImage. 211, 116631 (2020). https://doi.org/10.1016/j.neuroimage.2020.116631

